# A role for CIZ1, a Nuclear Matrix Protein Overexpressed in Cancer Cells, in Controlling the Initiation of DNA Replication

**DOI:** 10.64898/2026.10.10.758243

**Authors:** Manzar Hossain, Michael Tramantano, Sanjay Naik, Bruce Stillman

**Affiliations:** Cold Spring Harbor Laboratory, 1 Bungtown Road, Cold Spring Harbor, NY 11724; Department of Dermatology, The Ohio State University, Comprehensive Cancer Center, Columbus, OH 43210; Current Address

**Keywords:** CIZ1, Origin Recognition Complex (ORC), CDC6, Cyclin-dependent Kinase, DNA Replication

## Abstract

DNA replication licensing is tightly coordinated with cell-cycle progression to ensure genome stability. Here we demonstrate that CIZ1, a nuclear matrix scaffold protein in human cells, coordinates DNA replication licensing through direct interactions with the Origin Recognition Complex (ORC), CDC6, and Cyclins E/A–CDK2. The C-terminal zinc finger region of CIZ1 mediates binding to the cyclin-binding motifs of ORC1 and CDC6, and Cyclin–CDK-dependent phosphorylation regulates these interactions. Cell cycle analyses demonstrated a dynamic association of CIZ1 with ORC1 and CDC6. Functional studies showed that CIZ1 depletion impaired MCM loading, reduced DNA synthesis, delayed S-phase progression and inhibited cell proliferation. Cancer cells overexpressing CIZ1 exhibited an enhanced dependency on its expression, and CRISPR tiling screens identified the C-terminal regulatory region containing cyclin-binding motifs and CDK phosphorylation sites as essential for cell proliferation. Acute degradation of CIZ1 further revealed an unexpected role in maintaining CCCTC-binding factor (CTCF) occupancy at specific genomic loci and regulating transcriptional programs associated with cell adhesion and migration, indicating that CIZ1 links DNA replication with higher-order genome organization. Finally, depletion of CIZ1 significantly suppressed colorectal tumor growth in xenograft models. Together, these findings establish CIZ1 as a multifunctional scaffold that integrates DNA replication licensing, Cyclin– CDK signaling, and chromatin architecture.

## Introduction

The faithful and timely replication of the genome is a fundamental requirement for cell proliferation and the maintenance of genomic stability. In eukaryotes, DNA synthesis is initiated at defined chromosomal regions termed replication origins, whose activation is tightly regulated by the cell cycle machinery to guarantee that the genome is duplicated once, and only once, per cell division cycle (Costa and Diffley 2022; Hu and Stillman 2023; Frisbie and Bleichert 2026; You and Masai 2024; Stillman et al. 2025; Bleichert et al. 2017a). This temporal control is enforced through a two-step mechanism, the first of which is origin licensing by assembling a pre-Replicative Complex (pre-RC) at all potential origins of DNA replication across the entire genome during the G1 phase of the cell division cycle. Then, driven by the concerted action of CDC7-DBF4 (DDK) and Cyclin-dependent (CDK) kinases, each origin is activated in a temporal pattern throughout S phase. Dysregulation of either step is a hallmark of cancer, where aberrant replication initiation contributes to replication stress, DNA damage accumulation, and genomic instability that fuel tumor evolution (Macheret and Halazonetis 2015; Kotsantis et al. 2018; Igarashi et al. 2024). Unraveling the molecular mechanisms that link replication licensing to cell cycle progression is critical not only for advancing our understanding of fundamental cell biology, but also for uncovering therapeutic vulnerabilities that can be targeted in cancer.

The Origin Recognition Complex (ORC), a hetero-hexameric AAA+ ATPase comprising ORC1–6, serves as the master organizer of replication initiation by marking or licensing origins of DNA replication (Bleichert et al. 2017b; Attali et al. 2021). ORC recruits CDC6, an additional AAA+ ATPase, which together with CDT1 facilitates the loading of the MCM2–7 double hexamer, thereby licensing origins for replication (Parker et al. 2017; Weissmann et al. 2024; Evrin et al. 2009; Wells et al. 2025; Yang et al. 2024; Remus et al. 2009). The subsequent activation of licensed origins requires phosphorylation of MCM2–7 by the CDC7-DBF4 (DDK) kinase and the S-phase kinases Cyclin E–CDK2 and Cyclin A–CDK2, which also phosphorylate CDC6 and ORC1 to prevent re-licensing once S phase has commenced (Galanti and Pfander 2026; Li et al. 2023; Nasheuer and Meaney 2024; Palm and Costa 2025; Araki 2010). In human cells, ORC1 and CDC6 protein levels are themselves dynamically regulated across the cell cycle: ORC1 is degraded or displaced from chromatin during S phase and mitosis and is resynthesized in late G2 phase (Tatsumi et al. 2003; Ohta et al. 2003; Kara et al. 2015; Hossain et al. 2021), while CDC6 undergoes de novo synthesis during G1 phase following mitogenic stimulation (Petersen et al. 2000; Hossain et al. 2021). Tight temporal regulation of replication licensing factors, including ORC, CDT1 and CDC6, restricts DNA replication to a single round per cell cycle (Pozo and Cook 2016; Sugimoto et al. 2009; Hall et al. 2008; Liu et al. 2007; Coleman et al. 2015; Vaziri et al. 2003). In cancer cells, however, deregulated CDC6 expression and perturbations in ORC function are frequently associated with replication stress, genomic instability, and malignant progression (Yadav and Polasek-Sedlackova 2024; Igarashi et al. 2024).

Beyond the biochemical regulation of individual replication factors, the spatial organization of the replication machinery within the nucleus plays an equally important role in determining the temporal control of DNA replication, as well as the efficiency and fidelity of DNA replication. Early cell biological studies established that DNA replication occurs at discrete nuclear sites, termed replication foci, which are anchored to a proteinaceous nuclear scaffold or nuclear matrix (Anachkova et al. 2005; Berezney et al. 2000; Cook 1999). The nuclear matrix, operationally defined as the detergent-and nuclease-resistant nuclear fraction, provides a structural framework to which replication origins and associated proteins are tethered, thereby concentrating replication factors and facilitating the coordination of origin firing across the genome (Wilson and Coverley 2013; Djeliova et al. 2001). Several proteins have been implicated in linking replication origins to the nuclear matrix, but the molecular mechanisms governing this spatial organization remain incompletely understood, particularly in human cells.

CDKN1A-interacting zinc finger protein 1 (CIZ1) was originally identified as a binding partner of the CDK inhibitor p21^Cip1/Waf1^ and was subsequently shown to localize to the nuclear matrix, where it promotes DNA replication initiation in a cyclin–CDK-dependent manner (Coverley et al. 2005; Copeland et al. 2010; Mitsui et al. 1999). Studies in mouse cells demonstrated that Ciz1 is required for efficient S-phase entry and associates with nuclear replication foci, with its activity dependent on phosphorylation by Cyclin E– CDK2 (Copeland et al. 2015, 2010; Pauzaite et al. 2016; Munkley et al. 2011). The C-terminal domain of Ciz1, which contains three conserved zinc finger motifs and a matrin homology domain is necessary for nuclear matrix targeting (Ainscough et al. 2007). However, the molecular basis for CIZ1 interactions with core components of the pre-RC, particularly the ORC and CDC6, has remained unclear, and the functional characterization of human CIZ1 in DNA replication initiation has been limited. In addition to its replication-associated functions, CIZ1 has been reported to play roles in X chromosome inactivation maintenance (Ridings-Figueroa et al. 2017; Rodermund et al. 2021) and chromatin organization (Stewart et al. 2019), highlighting its multifunctional nature within the nucleus.

Mounting evidence implicates CIZ1 as a clinically relevant factor in human malignancy. Transcriptomic and proteomic analyses have identified CIZ1 overexpression in a range of solid tumours, including colorectal, lung, liver and breast cancers, where elevated CIZ1 levels correlate with advanced tumor stage and poor patient prognosis (Zhou et al. 2018; Wan et al. 2023; Wu et al. 2016; Wang et al. 2013; Chen et al. 2019; Zhang et al. 2015; Swarts et al. 2018; Turvey et al. 2025; Higgins et al. 2012). In colorectal cancer, CIZ1 has been reported to promote tumor growth and resistance to apoptosis, while CIZ1 depletion impairs cancer cell proliferation (YIN et al. 2013; Liu et al. 2016). Furthermore, recurrent mutations and splice variants of CIZ1 have been identified in leukemia and other hematological malignancies, suggesting that both gain-of-function and loss-of-function alterations may contribute to disease in different cellular contexts (Pauzaite et al. 2016; Liu et al. 2016; Swarts et al. 2018). Despite these associations, the mechanistic basis by which CIZ1 promotes cancer cell proliferation, and in particular whether its oncogenic activity is mechanistically linked to its replication scaffolding function, has not been established. Critically, whether CIZ1 overexpression creates a dependency in cancer cells has not been directly tested in vivo.

Here, we address these outstanding questions through a comprehensive biochemical, cell biological, and in vivo functional analysis of human CIZ1. We demonstrate that CIZ1 directly interacts with ORC1 and CDC6 through its C-terminal ZnF2/ZnF3 domain, engaging the Cy motifs of both replication factors in a manner regulated by Cyclin E–CDK2 and Cyclin A–CDK2. We show that ORC1 and CDC6 compete for binding to the same CIZ1 interface, revealing a novel regulatory mechanism by which the nuclear matrix coordinates pre-RC assembly and dissolution in a cell cycle-dependent fashion. In pursuing these experiments, significant differences in the protein interactions of human and mouse CIZ1 emerged. We further establish that CIZ1 is required for efficient pre-RC assembly and S-phase progression in human cells, and that tumor growth is dependent on CIZ1 in vivo. Using an inducible degron system for acute CIZ1 depletion, we further show that CIZ1 loss disrupts CTCF occupancy at a discrete subset of genomic sites and perturbs transcriptional programs governing cell motility, revealing an additional role for CIZ1 in CTCF-dependent genome organization beyond its function in replication licensing. Together, these findings identify CIZ1 as a critical scaffold integrating replication licensing with cyclin–CDK cell cycle control and nominate its functional interaction domain as a promising molecular target in CIZ1-driven cancers.

## Results

### Direct association of CIZ1 protein to ORC1 and CDC6 proteins links DNA replication to nuclear matrix

While investigating the proteins that interact with ORC and CDC6 in human cells, we uncovered a link with CIZ1 and because of previous studies showing a role for CIZ1 in DNA replication (Copeland et al. 2010; Coverley et al. 2005), we further investigated these protein-protein interactions. Whole cell lysate and chromatin extract prepared from asynchronously proliferating U2OS cells were used for CIZ1 immunoprecipitation using two different antibodies. Immunoprecipitation results revealed that CIZ1 interacted with several DNA replication proteins such as ORC1, ORC3 and CDC6 (Fig. 1*A* and Fig. S1*A*). An association between CIZ1 and CDC6 has been reported (Copeland et al. 2015). Surprisingly, a previously been reported interaction between p21^Cip1/Waf1^ and CIZ1 protein in the human cells was not observed (Mitsui et al. 1999). The spatial-temporal localization and expression of CIZ1 protein during the human cell cycle were further studied. In the U2OS cell line, mGFP was fused to the CIZ1 protein, with expression controlled by a tetracycline regulated CMV promoter and GFP localization was followed upon the addition of tetracycline. The mGFP-CIZ1 showed a typical nuclear matrix localization with discrete dot-like subnuclear pattern throughout the nucleus in interphase cells. Interestingly, the mGFP-CIZ1 protein was absent from mitotic chromosomes upon nuclear envelope breakdown, but and then redecorated the chromosomes in the newly emerged daughter cells (Fig. 1B). Upon transient transfection of mCherry-CIZ1 into a U2OS cell that stably expressed mGFP-ORC1, both proteins partially co-localized in the nucleus with or without Triton-X-100 extraction (Fig. 1C). Because of dynamic spatial distribution of human CIZ1 proteins we examined the temporal expression of the protein during the cell division cycle. HeLa suspension cells were synchronized with a double thymidine block and released and then CIZ1 and other proteins were detected by immunoblot (Fig. 1D). CIZ1 levels fluctuated slightly during the cell cycle, with the highest levels during late G2/M and the mobility of CIZ1 protein was retarded during the mitotic phase, possibly due to phosphorylation which has been observed (Copeland et al. 2015).

**Fig. 1.**
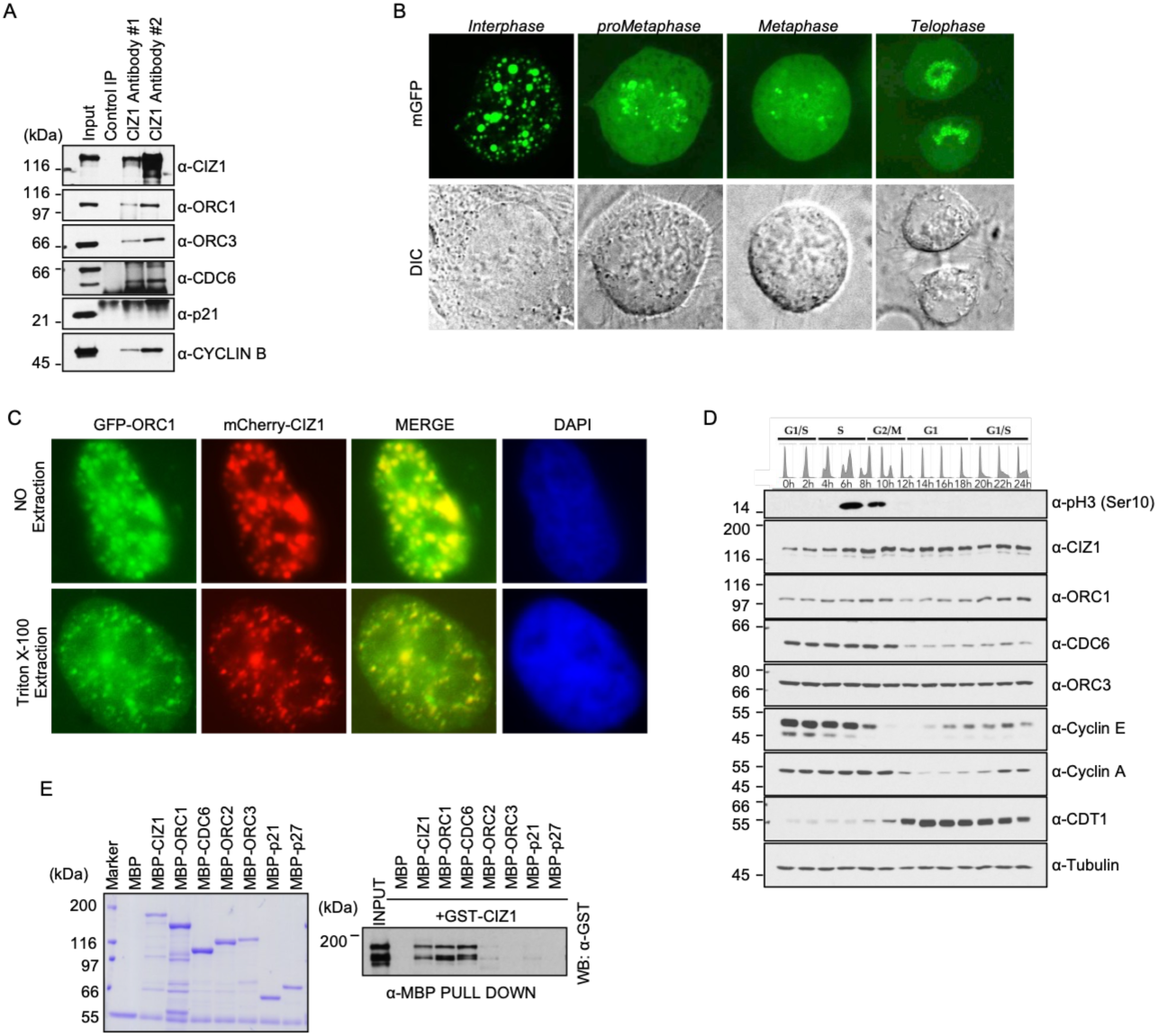
CIZ1 is cell cycle regulated and interacts with pre-RC components. (A) Immunoprecipitation of CIZ1 from asynchronously proliferating U2OS chromatin lysates showing interactions with ORC1, CDC6, ORC3 and Cyclin B. Input and IP are 5% and 10%, respectively. Molecular weight markers, kDa. (B) Representative images of tetracycline-inducible mGFP-CIZ1 U2OS cells in interphase and mitosis. The bottom panels show the corresponding differential interference contrast (DIC) images. (C) Tetracycline-inducible GFP-ORC1 stable U2OS cells were transiently transfected with mCherry-CIZ1. Protein co-localization was examined in cells with or without Triton X-100 extraction. Images were acquired at 40× magnification. (D) Expression profiles of CIZ1, the pre-RC proteins ORC1, ORC3, CDC6, and CDT1, and the cell-cycle regulated proteins Cyclin E, Cyclin A, and phospho-histone H3 (Ser10) were examined by immunoblotting in double-thymidine-block synchronized HeLa suspension cells following release. DNA content was analyzed by flow cytometry at the indicated time points (hours) after release, with the corresponding cell-cycle phases indicated. (E) Purified MBP-CIZ1, MBP-ORC1, MBP-ORC3, MBP-CDC6, MBP-p21^CIP1/WAF1^ and MBP-p27^KIP1^ protein bound to α-MBP antibody magnetic beads were incubated with purified GST-CIZ1 protein in an MBP pull-down assay and blotted with anti-GST antibody (bottom panel). The loading of MBP-fused recombinant proteins used in the pull-down assay is shown in top panel. MBP protein served as negative control.

To further investigate the type of interaction between human CIZ1 protein and DNA replication protein, recombinant proteins expressed as Maltose Binding Protein (MBP) or Glutathione S-transferase (GST) fusion proteins were expressed in bacteria and purified. In MBP immunoprecipitation assays using purified proteins, GST-CIZ1 interacted with MBP-CIZ1, MBP-ORC1 and MBP-CDC6, while no direct interaction was detected with other ORC subunits such as MBP-ORC2 and MBP-ORC3 (Fig. 1E). Consistent with the in vivo results, a direct interaction between GST-CIZ1 and either MBP-p21^CIP1/WAF1^ or MBP-p27^KIP1^ proteins was not observed (Fig. 1E). A direct interaction between CIZ1 and ORC1, but not the other ORC subunits, was observed (Fig. S1B).

### Comparative analysis of Human and Mouse CIZ1 Protein Interactions

The lack of interaction between human CIZ and p21 ^Cip1/Waf1^ was puzzling given the report of 2-hybrid interactions in yeast (Mitsui et al. 1999). We therefore investigated the interaction between p21 ^Cip1/Waf1^ and CIZ1 proteins from both human and mouse. Sequence analysis and domain architecture of the human CIZ1 protein reveal strong resemblance to its mouse counterpart (Fig. 2A). The full-length human and mouse CIZ1 have approximately 70% overall homology at the amino-acid level. The N-terminus of human CIZ1 protein is mostly disordered while its C-terminus is predicted to be more structured (Fig. 2B). As shown in the schematic, both proteins share a similar domain architecture, with the N-terminus harboring two well-separated glutamic acid-rich regions (polyQ), while the C-terminus contains the major structured domains (amino acids 597-730; 780-832 in human). The C-terminal region of both human and mouse CIZ1 proteins contain three highly conserved zinc finger domains, an acidic domain, and a matrix-anchoring Matrin homology domain toward the C-terminus. For the immunoprecipitation assays, the mouse and human CIZ1 proteins, DNA replication proteins (ORC1, CDC6, ORC2, and ORC3), and CDK kinase inhibitors (p21^Cip1/Waf1^ and p27^Kip1^) were expressed and purified from bacterial cells as MBP- or GST-fusion proteins. The binding studies demonstrated that both human and mouse CIZ1 proteins directly interacted with the DNA replication proteins ORC1 and CDC6. In addition, mouse Ciz1 showed an extra interaction with ORC2, whereas no interaction was observed with ORC3. Interestingly, mouse Ciz1 directly interacted with p21^WAF1/CIP1^ but not with p27^Kip1^, whereas human CIZ1 did not show interaction with either protein, consistent with observations described above (Fig. 2C). Human CIZ1 protein interacted with ORC1 and CDC6 proteins through the C-terminal 550-898 amino acid region containing the structured domains (Fig. 2D). Similarly, the mouse Ciz1 protein also used the same C-terminal domain to interact with mouse ORC1 and CDC6 proteins (Fig. 2D and Fig. S2A), in contrast to previous findings about CDC6 (Copeland et al. 2010; Ainscough et al. 2007; Copeland et al. 2015). Further investigation of protein-protein interactions showed that the region from CIZ1 655-720 aa, containing the ZnF2 and ZnF3 domains, was sufficient for interaction with human ORC1 and CDC6 proteins (Fig. 2E and Figs. S2B, S2C S2D and S2E). Internal deletion of the structured do mains within the C-terminal half of the human CIZ1 protein indicated deletion of both zinc fingers ZnF2 and ZnF3 compromised ORC1 interaction, whereas deletion of the ZnF2+ZnF3 plus amino acids more C-terminal to these Zn fingers was required to compromise CDC6 interaction (Fig. 2F and Figs. S2F and S2G).

**Fig. 2.**
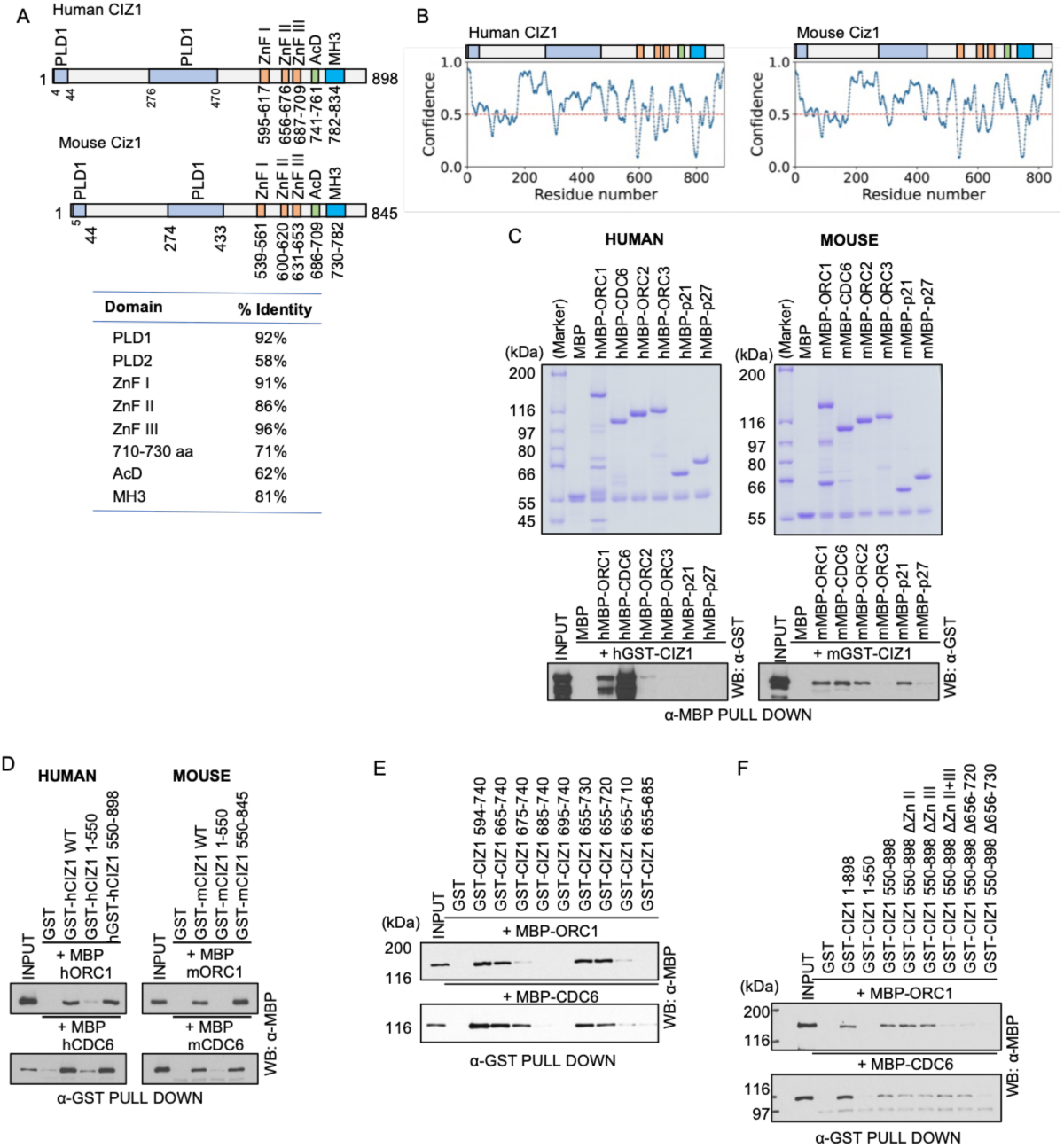
Direct protein-protein interaction of CIZ1 with DNA Replication proteins is conserved. (A) Schematic showing domain architecture of human CIZ1 and mouse Ciz1 proteins. Bottom table showing primary sequence identity between different domains of the human and murine proteins. PLD, Prion-like domain; ZnF, Zinc-finger domain; AcD, glutamic and aspartic amino acid-rich acidic domain; MH3, Matrin 3 Homology 3. (B) Prediction of intrinsic disorder profile of human CIZ1 and mouse Ciz1 proteins with PrDOS. The C-terminal half of the proteins is more structured compared to the N-terminus. (C) Purified MBP-ORC subunit proteins ( ORC1, ORC2 and ORC3), MBP-CDC6 protein and CDK inhibitors (MBP-p21^CIP1/WAF1^ and MBP-p27^KIP1^) bound to anti-MBP antibody magnetic beads were incubated with purified GST-CIZ1 protein in an MBP pull-down assay and blotted with anti-GST antibody (bottom panel). The loading of MBP-fused recombinant proteins used in the pull-down assay is shown in top panel. MBP protein served as a negative control. Left panel, human protein interactions; Right panel, mouse protein interactions. (D) Equimolar amounts of GST-CIZ1 wild type as well as its N- and C-truncation mutants (human, 1-550 and 550-898; mouse, 1-550 and 550-845) were incubated with either full-length MBP-ORC1 (top panel) or MBP-CDC6 (bottom panel) proteins in a GST antibody-based binding assay and blotted with anti-MBP antibody. Left panel, human proteins; Right panel, mouse proteins. *See also Figure S2A*. (E) The C-terminal deletion mutants of human GST-CIZ1 protein were bound to α-GST antibody magnetic beads were incubated either with purified MBP-ORC1 (top panel) or MBP-CDC6 (bottom panel) proteins in a GST pull-down assay and blotted with anti-MBP antibody. *See Figure S2D and S2E*. (F) The full length, N-terminus, C-terminus and various internal deletion mutants of GST-CIZ1 recombinant proteins pre-bound to magnetic beads were incubated either with purified MBP-ORC1 or MBP-CDC6 proteins. The interaction was analyzed by blotting with anti-MBP antibody (top panel, MBP-ORC1; bottom panel, MBP-CDC6). *See Figure S2F and S2G*.

### Shared binding region of CIZ1 to ORC1 and CDC6 reveal a regulatory mechanism for DNA replication

Since the ORC1 and CDC6 binding regions overlap within the CIZ1 protein, we investigated the CIZ1 binding regions within these proteins. To map the CIZ1 binding region within human ORC1 and CDC6 proteins the full-length GST-CIZ1 protein was incubated with series of N- or C-terminal fragments of MBP-ORC1 and MBP-CDC6 in MBP pull down assays. The results indicated that the human ORC1 protein required its N-terminal region in making contacts with CIZ1 protein (Fig. 3A and Fig, S3A). Interestingly, this intrinsically disordered region (IDR) of ORC1 contains the amino acids in that bind to Cyclin A and CDC6 (see Fig. 3D and ref (Hossain et al. 2021)). We therefore tested mutations that affect Cyclin A or CDC6 interaction with ORC1 to determine if they also affect CIZ1 interaction. Mutation of five basic residues within the 1-300 aa region of MBP-ORC1, the M5A mutant that prevents both Cyclin A and CDC6 binding to ORC1, also compromised CIZ1 binding to ORC1. The M5A mutation in the full-length MBP-ORC1 protein substantially reduced its interaction with GST-CIZ1, while other previously described basic patch mutants BP4 and BP5 that regulate the binding of CDC6 to ORC1 (Hossain et al. 2021) still bound to CIZ1 (Fig. 3B and Fig, S3B). To further investigate this interaction, a series of single- and double-point mutations within part of the ORC1 IDR fragment (100-250) targeting the ORC1 Cy motif (Fig. 3D) (Hossain et al. 2021) were assessed for CIZ1 binding (Fig. 3C and Fig. S3C). Mutations within the core of the ORC1 Cy motif (ORC1²³⁵KRL²³⁷ to ORC1 ³⁵ARA²³⁷) disrupted ORC1-CIZ1 binding, whereas point mutations flanking the core Cy motif displayed differential binding to CIZ1 (Fig. 3C and Fig. S3C). Notably, mutation of L237 as well as residues flanking the core Cy motif, specifically TP230/231 and EL238/239, were dispensable for CIZ1 binding (Fig. 3C and Fig. S3C).

**Fig. 3.**
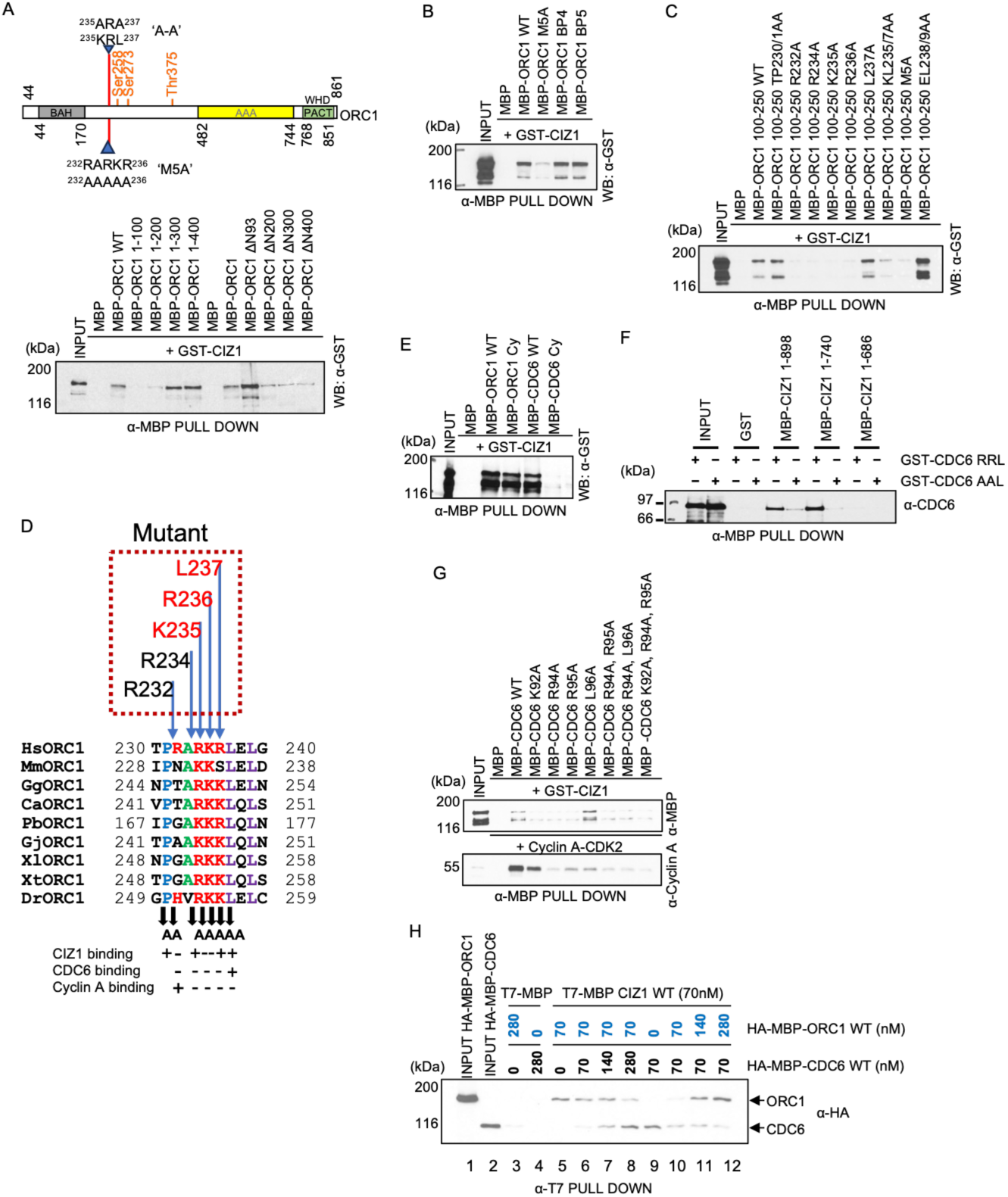
Human ORC1 competes with CDC6 protein for binding to CIZ1. (A) Schematic of ORC1 domain architecture (BAH, AAA+ ATPase, PACT, Winged Helix (WH) domains) highlighting the region mutated to generate the ’A-A’ (^235^ARA^237^) and ’M5A’ (^232^AAAAA^236^) mutants, and phosphorylation sites (Ser258, Ser273, Thr375). Below, MBP pull-down assays using MBP-ORC1 WT and a series of N-terminal truncations/deletions (1–100 through ΔN400) incubated with GST-CIZ1, immunoblotted with anti-GST. (B) The anti-MBP pull-down of MBP, MBP-ORC1 1–861 WT, MBP-ORC1 M5A, MBP-ORC1 BP4 or MBP-ORC1 BP5 with GST-CIZ1 and immunoblotted with anti-GST antibody. (C) The anti-MBP pull-down of MBP-ORC1 100–250 WT and a panel of point/cluster mutants (TP230/231AA, R232A, R234A, K235A, R236A, L237A, KL235/237AA, M5A, EL238/239AA) with GST-CIZ1 and immunoblotted with anti-GST antibody. (D) Multiple sequence alignment of the ORC1 motif across species (human, mouse, chicken, dog, opossum, zebra finch, frog, zebrafish), with mutated residues indicated above and a summary table below showing mutations that differentially affect CIZ1, CDC6, and Cyclin A binding. (E) The anti-MBP pull-down of MBP, MBP-ORC1 WT, MBP-ORC1 Cy mutant [KL235/237AA], MBP-CDC6 WT or MBP-CDC6 Cy mutant [CDC6^94^RRL^96^ to CDC6^94^AAL^96^] with GST-CIZ1 WT, immunoblotted with anti-GST antibody. (F) The anti-MBP pull-down of MBP, MBP-CIZ1 WT^1-898^, MBP-CIZ1-^1-740^ or MBP-CIZ1^1-686^ with GST-CDC6 WT or GST-CDC6 Cy mutant [CDC6^94^RRL^96^ to CDC6^94^AAL^96^], immunoblotted with anti-CDC6 antibody. (G) The anti-MBP pull-down of MBP, MBP-CDC6 WT and MBP-CDC6 mutants (K92A, R94A, R95A, L96A, and combinations) with either GST-CIZ1 (top, anti-GST) or Cyclin A-CDK2 (bottom, anti-Cyclin A), identifying CDC6 residues required for CIZ1 versus Cyclin A binding. (H), The anti-T7 of MBP-T7 or T7-MBP-CIZ1 WT pull-down with various concentrations of HA-MBP-ORC1 and HA-MBP-CDC6 (0–280 nM), immunoblotted with anti-HA.

Since CIZ1 bound to the ORC1 Cy-motif region, we investigated whether the CDC6 Cy motif, CDC6^94^RRL^96^, that is required for its interaction human Cyclin E, Cyclin A and ORC1 (Hossain et al. 2021; Hossain and Stillman 2016) was also involved in interaction with CIZ1 protein. Mutation of the core residues in the CDC6 Cy motif [CDC6^94^RRL^96^ to CDC6^94^AAL^96^, see Refs (Hossain et al. 2021; Hossain and Stillman 2016)] completely abolished its interaction with CIZ1 protein (Fig. 3E and Fig. S3D). The results were further confirmed using different MBP-CIZ1 truncation mutants that showed defective binding with the CDC6 Cy mutant (Fig. 3F and Fig S3E). To determine the critical residues near the CDC6 Cy motif involved in its interaction with CIZ1 protein, a series of point mutations within recombinant full-length MBP-CDC6 protein were tested for binding to both Cyclin A-CDK2 and GST-CIZ1. The results of multiple mutants showed that the CIZ1 binding domain in CDC6 overlapped with the CDC6 Cy motif that binds Cyclin A, but they were clearly distinguishable (Fig. 3G and Fig. S3F and S3G). Thus, the Cyclin A binding regions of ORC1 and CDC6 were required for binding to CIZ1.

Since both ORC1 and CDC6 interact with each other as well as their interacting regions on CIZ1 are very close and overlapping, we investigated whether the replication proteins were cooperative or competitive for binding the Zinc finger domains of CIZ1, thereby regulating the location of replication on the nuclear matrix. First, an in vitro competition assay was performed with purified T7-MBP-CIZ1, HA-MBP-ORC1 and HA-MBP-CDC6 proteins. An increasing concentration of HA-MBP-ORC1 inhibited the interaction between HA-MBP-CDC6 and T7-MBP-CIZ1 in a stoichiometric ratio (Fig. 3H, lanes 9-12). In contrast, increasing concentrations of HA-MBP-CDC6 protein had a limited effect on the interaction between HA-MBP-ORC1 and CIZ1-ORC1 (Fig. 3H, lanes 5-8). In conclusion, these findings demonstrate that the ORC1 and CDC6 Cy motifs play critical roles in mediating their interaction with CIZ1, distinct from the interaction with Cyclin A. Furthermore, the competitive binding observed between ORC1 and CDC6 for CIZ1 suggests a regulatory mechanism in which these replication factors dynamically coordinate association with CIZ1 to control replication complex positioning on the nuclear matrix.

### Regulation of CIZ1–ORC1 and CIZ1–CDC6 interactions by Cyclin E- and Cyclin A-dependent CDK activity

Cyclin-CDK–dependent phosphorylation of human CIZ1 was investigated. In vitro kinase assays demonstrated that the C-terminal fragment of CIZ1 (amino acids 550–898) but not the amino-terminal fragment (amino acids 1-550) was phosphorylated by both Cyclin E-CDK2 and Cyclin A-CDK2 kinase complexes (Fig S4A). This was surprising because Cyclin-dependent protein kinase phosphorylation of mouse CIZ1 has been studied, showing phosphorylation in the amino-terminal region (Copeland et al. 2015; Coverley et al. 2005). We therefore examined the phosphorylation of the N- and C-terminal fragments of mouse CIZ1, showing that Cyclin E-CDK2 phosphorylated both fragments and Cyclin A-CDK2 phosphorylated primarily the C-terminal fragment (Fig. S4B). Like the differential binding to p21 ^Cip1/Waf1^, these phosphorylation studies uncovered another difference between the mouse and human CIZ1 proteins.

The involvement of the Cy motifs of the DNA replication proteins ORC1 and CDC6 in mediating their interaction with CIZ1 prompted investigation of the presence of Cyclin-binding motifs and Cyclin–CDK phosphorylation sites within human CIZ1. Detailed sequence analysis of human CIZ1 protein identified three potential Cyclin-binding RxL motifs within its C-terminal region, suggesting their possible involvement in binding to the G1/S phase Cyclin–CDK complexes, Cyclin E–CDK2 and Cyclin A–CDK2. Notably, the tandem Cy1 and Cy2 motifs are located in close proximity to the zinc finger 3 domain of CIZ1, whereas the third Cy3 motif is positioned at the extreme C-terminus of the protein (Figs. 2A and 4A). To test the involvement of in silico found Cy motifs of CIZ1 in making contact with the Cyclin-CDK complexes, a series of single, double and triple Cy motif mutants were made in CIZ1 and purified from bacteria as GST-fusion proteins. Binding studies demonstrated that all the three Cy motifs interacted with Cyclin E-CDK2 and Cyclin A-CDK2 and only when all three were mutated was binding compromised (Fig. 4B and Figs. S4C and S4D). However, we did not observe any reduction in phosphorylation in the Cy motif mutants of either full length or the C-terminal fragment of human CIZ1 protein (Fig. S4A and S4E). Furthermore, in silico analysis identified three canonical CDK consensus phosphorylation motifs (S/TPxR/K) within the C-terminal region of CIZ1, corresponding to threonine residues at positions T585, T686, and T876 (Fig. 4C). GST-CIZ1 wild-type and phosphorylation-site single mutants (T585A, T696A, T876A) and double/triple combinations were subjected to in vitro kinase assays using either Cyclin E-CDK2 or Cyclin A-CDK2. With Cyclin E-CDK2, wild-type CIZ1 and single-site mutants retain substantial phosphorylation, whereas combined T585A/T876A and triple mutants show progressively reduced signal, indicating that T585 and T876 were the predominant Cyclin E-CDK2 phosphorylation sites (Fig. 4D). In contrast, Cyclin A-CDK2 phosphorylated wild-type CIZ1 robustly, and the signal was only modestly reduced for T585A/T876A and triple mutants, suggesting broader or redundant phosphorylation site usage by Cyclin A-CDK2 (Fig. 4D).

**Fig. 4.**
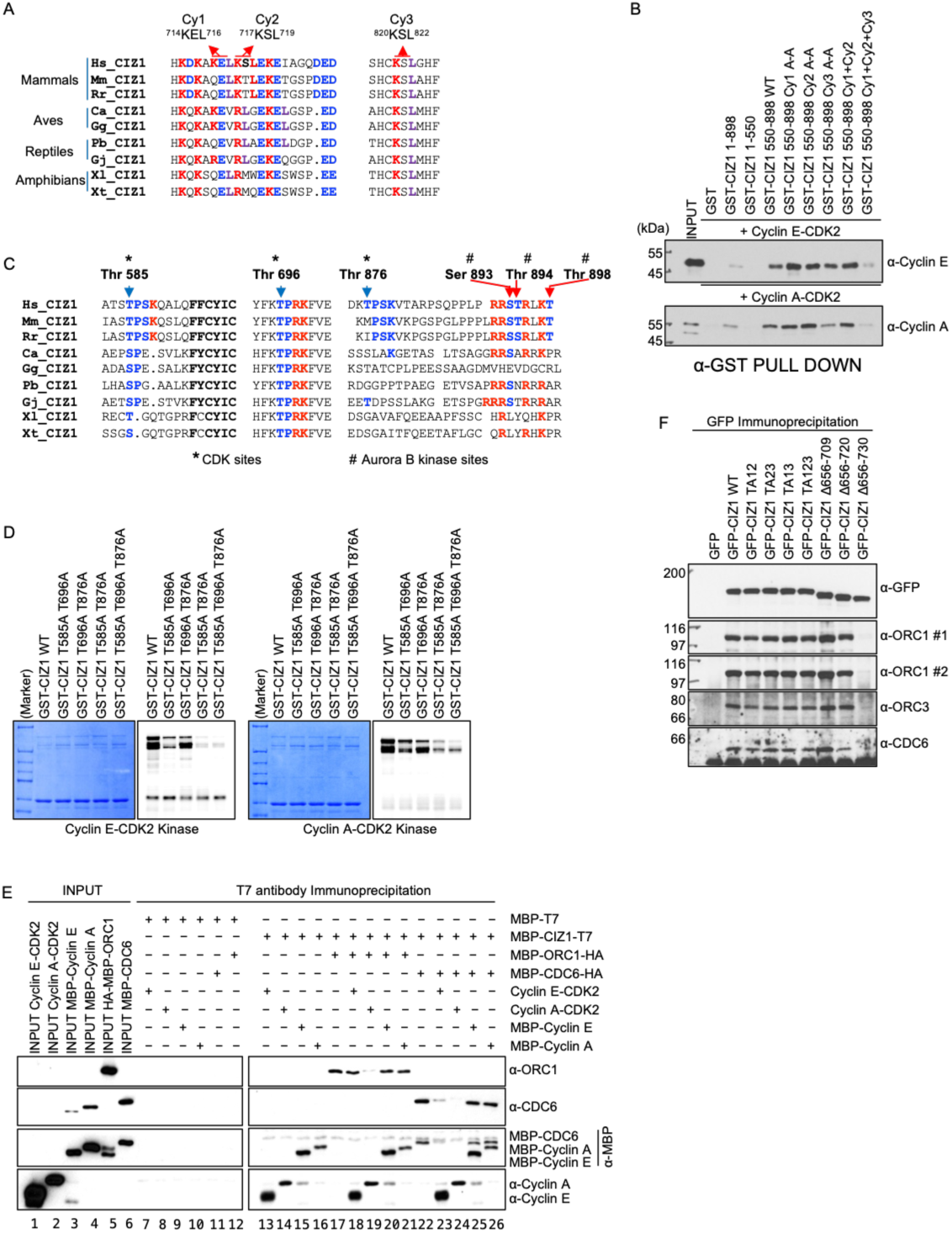
CIZ1 harbors conserved Cyclin-binding (Cy) motifs and CDK phosphorylation sites that regulate its interaction with ORC1, CDC6, and Cyclin-CDK2 complexes. (A) Multiple sequence alignment of ORC1 across mammals, birds, reptiles, and amphibians, highlighting three conserved potential cyclin-binding (Cy) motifs: Cy1 (^714^KEL^716^), Cy2 (^717^KSL^719^), and Cy3 (^820^KSL^822^). (B) GST pull-down of GST-CIZ1 1–898, 1–550, and 550–898 WT or Cy-motif mutants (Cy1 A-A, Cy2 A-A, Cy3 A-A, Cy1+Cy2, Cy1+Cy2+Cy3) incubated with either Cyclin E-CDK2 or Cyclin A-CDK2, immunoblotted with anti-Cyclin E and anti-Cyclin A. (C) Multiple protein sequence alignment of CIZ1 across species showing three conserved potential CDK phosphorylation sites (Thr585, Thr696, Thr876, marked with asterisks) and three Aurora B kinase sites (Ser893, Thr894, Thr898, marked with #). (D) In vitro kinase assays with Cyclin E-CDK2 (left) or Cyclin A-CDK2 (right) on GST-CIZ1 WT or phospho-site mutants (T585A/T696A/T876A combinations). Coomassie-stained gels (input) and corresponding autoradiographs/phosphor-images show that combined mutation of these threonines progressively reduced CIZ1 phosphorylation by both Cyclin-CDK2 complexes. (E) In vitro reconstitution assay using MBP-T7, MBP-CIZ1-T7, MBP-ORC1-HA, MBP-CDC6-HA, Cyclin E-CDK2, Cyclin A-CDK2, MBP-Cyclin E or MBP-Cyclin A in the presence of ATP. Following T7 antibody immunoprecipitation, immunoblotting for ORC1, CDC6, MBP, and Cyclins showed phosphorylation affected binding of CIZ1 with ORC1, CDC6 in a multi-protein assembly. (F) GFP immunoprecipitation from cells expressing GFP alone, GFP-CIZ1 WT, phospho-site mutants (TA12, TA23, TA13, TA123), or internal deletion mutants (Δ656–709, Δ656–720, Δ656–730), immunoblotted for GFP, ORC1 (two antibodies), ORC3, and CDC6. Deletion mutants spanning residues 656–730 showed reduced co-precipitation of ORC1 and CDC6, while phospho-site mutations had minimal effect, confirming this region as critical for pre-RC component binding independent of CDK phosphorylation status.

Next, we sought to determine how Cyclin-CDK–mediated phosphorylation of CIZ1 regulates its interactions with ORC1 and CDC6. Co-immunoprecipitation (Co-IP) experiments were performed using a T7-tag antibody to investigate the protein-protein interactions between CIZ1 and key DNA replication initiation factors, specifically ORC1, CDC6 in presence of Cyclin E-CDK2 and Cyclin A-CDK2. In this assay, MBP-CIZ1-T7 served as the bait protein, while various combinations of MBP-ORC1-HA, MBP-CDC6-HA, Cyclin E-CDK2, Cyclin A-CDK2, MBP-Cyclin E, and MBP-Cyclin A were used as prey proteins. The ORC1 blot revealed that MBP-ORC1-HA was successfully pulled down by MBP-CIZ1-T7 across multiple conditions except in the presence Cyclin A-CDK2 or Cyclin E-CDK2, demonstrating that CIZ1 binds ORC1 directly and that this interaction is inhibited by phosphorylation by Cyclin A-CDK2 activity, but not by Cyclin E-CDK2 activity (Fig 4E, Lanes 17-19). Similarly, the CDC6 blot and MBP blot shows that MBP-CDC6-HA was co-immunoprecipitated with MBP-CIZ1-T7, confirming a direct physical association between CIZ1 and CDC6, while its interaction was greatly reduced (Cyclin E-CDK2) or completely abrogated (Cyclin A-CDK2) (Fig. 4E, lanes 22-24). The MBP blot also shows that MBP-CIZ1-T7 interacted with both MBP-Cyclin A and MBP-Cyclin E without having an effect on CIZ1 interaction with either ORC1 or CDC6 in their presence, ruling out the possibly of Cyclin competition in diminishing the interaction (Fig. 4E, lanes 20, 21, 25, 26). In a physiological context, anti-GFP immunoprecipitation from 293 cells transiently expressing GFP-CIZ1 wild-type or phosphorylation mutants (TA12, TA23, TA13, or TA123) co-precipitated ORC1, ORC3, and CDC6, as confirmed using two independent α-ORC1 antibodies. These interactions were preserved across all phospho-mutant and deletion constructs (Δ656– 709 and Δ656–720), indicating that binding of CIZ1 to ORC1, ORC3, and CDC6 is independent of its phosphorylation status. In contrast, the CIZ1 Δ656–730 mutant exhibited a complete loss of interaction with ORC1, ORC3, and CDC6 (Fig. 4F and Fig, S4F). Notably, CIZ1 Cy motif triple mutants, which are defective in Cyclin binding but not completely impaired in phosphorylation, also failed to associate with ORC1 and CDC6 in both in vitro and physiological contexts (Fig. S4G). These findings suggest that Cyclin-binding interfaces, rather than promoting phosphorylation per se, may be critical for CIZ1 \interaction with replication licensing factors, warranting further investigation. Taken together, these results establish that CIZ1 is capable of dynamically interacting with ORC1, CDC6, Cyclin E-CDK2, and Cyclin A-CDK2, positioning it as a central scaffolding protein at the nexus of replication initiation and cell cycle regulation. This multi-protein interaction capability suggests that CIZ1 plays a critical role in coordinating the assembly or stabilization of the pre-replication complex (pre-RC) at DNA replication origins, potentially facilitating the phosphorylation of licensing factors by Cyclin-CDK complexes to trigger the transition from a licensed to an active replication origin. The ability of CIZ1 to bridge both early-S-phase kinase activity (Cyclin E-CDK2) and late-S-phase kinase activity (Cyclin A-CDK2) with core ORC and the licensing machinery further implies that CIZ1 may function throughout S-phase progression, ensuring the timely and ordered firing of replication origins. Overall, this result provides compelling biochemical evidence that CIZ1 acts as a molecular platform integrating cell cycle signals with the DNA replication machinery, consistent with its proposed role in promoting efficient DNA replication initiation during S-phase entry.

### Integration of CIZ1 with ORC1 and CDC6 in the control of DNA replication initiation in human cells

Given the dynamic expression patterns of ORC1 and CDC6 across the cell cycle, we investigated their biochemical interactions with CIZ1. HeLa suspension cells were synchronized using a double thymidine block, and following release, cell extracts were prepared for co-immunoprecipitation. The association between CIZ1 and key DNA replication proteins (ORC1, ORC3, ORC4, CDC6, and MCM2) as well as Cyclin A were monitored by CIZ1 immunoprecipitation from HeLa cell lysates (Fig. 5A). CIZ1 associated with ORC1, CDC6, and Cyclin A during early mitosis, approximately 10 hours after the second thymidine release, followed by a concomitant reduction in these interactions. Upon entry into early G1 phase, CIZ1 progressively re-established its interaction with ORC1, and subsequently, in mid-G1, also regained its association with CDC6. CIZ1 interaction with ORC1 immediately after G2/M phase, followed by recruitment of CDC6 occurred at the time of CIZ1 association with MCM2. MCM2 increased its association with CIZ during G1 phase and remained associated throughout S phase and then was reduced as cells entered into late S and G2 phases. During S phase, MCM2 appeared as a double, possibly due to phosphorylation.

**Fig. 5.**
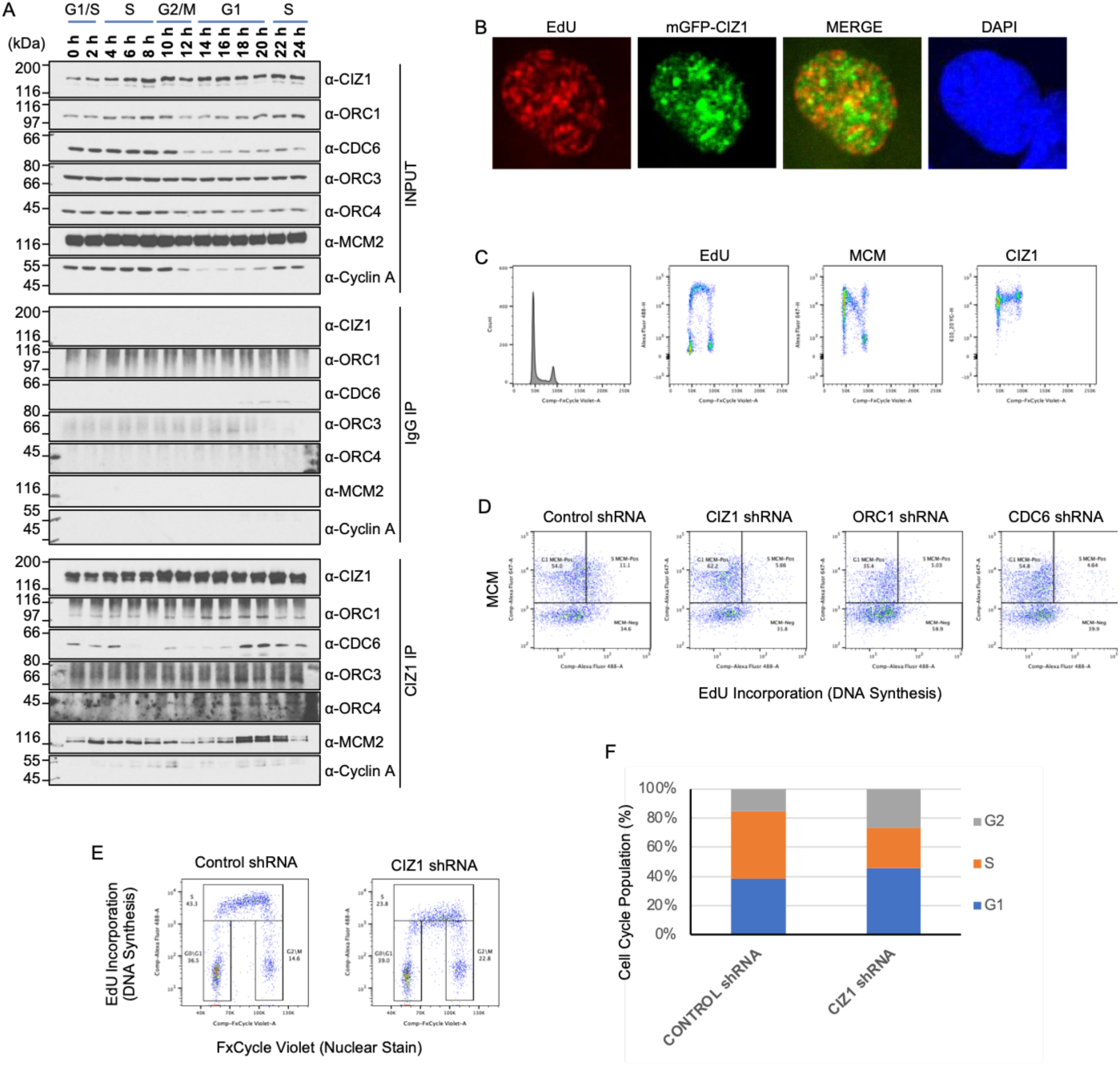
Dynamic association of CIZ1 with the pre-replication complex throughout the cell cycle and is required for efficient S-phase entry and DNA replication. (A) Synchronized HeLa cells were synchronized with a double thymidine block and released and harvested at the indicated time points spanning G1/S, S, G2/M, and G1 phases (0–24 h). Input lysates and CIZ1 or IgG immunoprecipitates were immunoblotted for CIZ1, ORC1, CDC6, ORC3, ORC4, MCM2, and Cyclin A. (B) Representative immunofluorescence images of EdU-labeled cells expressing mGFP-CIZ1, showing EdU (red, sites of active DNA synthesis), mGFP-CIZ1 (green), merged overlay, and DAPI (nuclear DNA). (C) Flow cytometry analysis of cell-cycle distribution (FxCycle Violet, left) and single-parameter plots of EdU incorporation, MCM loading, and CIZ1 levels across the cell cycle. (D) Flow cytometric analysis of MCM loading versus EdU incorporation in cells expressing control, CIZ1, ORC1, or CDC6 shRNA. Quadrant percentages indicate the proportion of cells in each category (G1 MCM-positive, S-phase MCM-positive, MCM-negative). CIZ1, ORC1, and CDC6 depletion each reduced the fraction of cells entering S phase (MCM-positive/EdU-positive) relative to control. (E) Representative cell-cycle profiles (EdU incorporation vs. FxCycle Violet nuclear stain) in control shRNA and CIZ1 shRNA cells, with gated percentages for G0/G1, S, and G2/M populations. (F) Quantification of cell-cycle distribution from (E), showing a reduced S-phase fraction and increased G1 and G2 fractions upon CIZ1 knockdown compared to control shRNA, consistent with a role for CIZ1 in promoting efficient S-phase progression.

To further examine the physiological significance of CIZ1 association with the DNA replication machinery at the G1/S boundary, we utilized a stable U2OS cell line expressing mGFP-CIZ1 and labeled it for 60 minutes with 5-ethynyl-2’-deoxyuridine (EdU) followed by immunofluorescence, which revealed limited co-localization of CIZ1 with sites of active DNA replication (Fig. 5B). This is in contrast to claims of co-localization of CIZ1 with sites of DNA replication (Coverley et al. 2005). Next, the association of CIZ1 with chromatin in HCT116 cells during the cell division cycle was assessed by 2-dimensional flow cytometry. Cells were labelled with EdU for 60 minutes, extracted with triton-X-100 to remove non-chromatin bound proteins and then stained with FxCycle Violet for DNA content. 2-dimsenional flow cytometry of singlet cells revealed distinct G1, S and G2/M populations (Fig 5C, EdU). Chromatin bound MCM, detected using anti-MCM2 antibody (Alexa Fluor 647), was observed in late G1 phase and decreased as cells progressed through S phase as previously observed (Moreno et al. 2016; Håland et al. 2015). In contrast, CIZ1 (610_20 YG) was detected in late G1 phase cells through to late S phase cells (Fig. 5*C*).

Depletion of CIZ1, ORC1, or CDC6 by doxycycline induced shRNA in HCT116 cells reduced the S-phase, MCM-positive population compared to control shRNA (11.1%), with CIZ1 shRNA (5.66%), ORC1 shRNA (5.03%), and CDC6 shRNA (4.64%), all showing approximately 50% reduction in MCM levels during S phase, indicating that CIZ1, like ORC1 and CDC6, is required for efficient DNA replication initiation (Fig. 5D, see Fig. 6B for knockdown efficiency). CIZ1 depletion by doxycycline-induced shRNA in HCT116 cells greatly reduced the S-phase population from 43.3% to 23.8%, with a concomitant increase in G2/M cells from 14.6% to 22.8%, while G0/G1 remained largely unchanged (Fig. 5E).

**Fig. 6.**
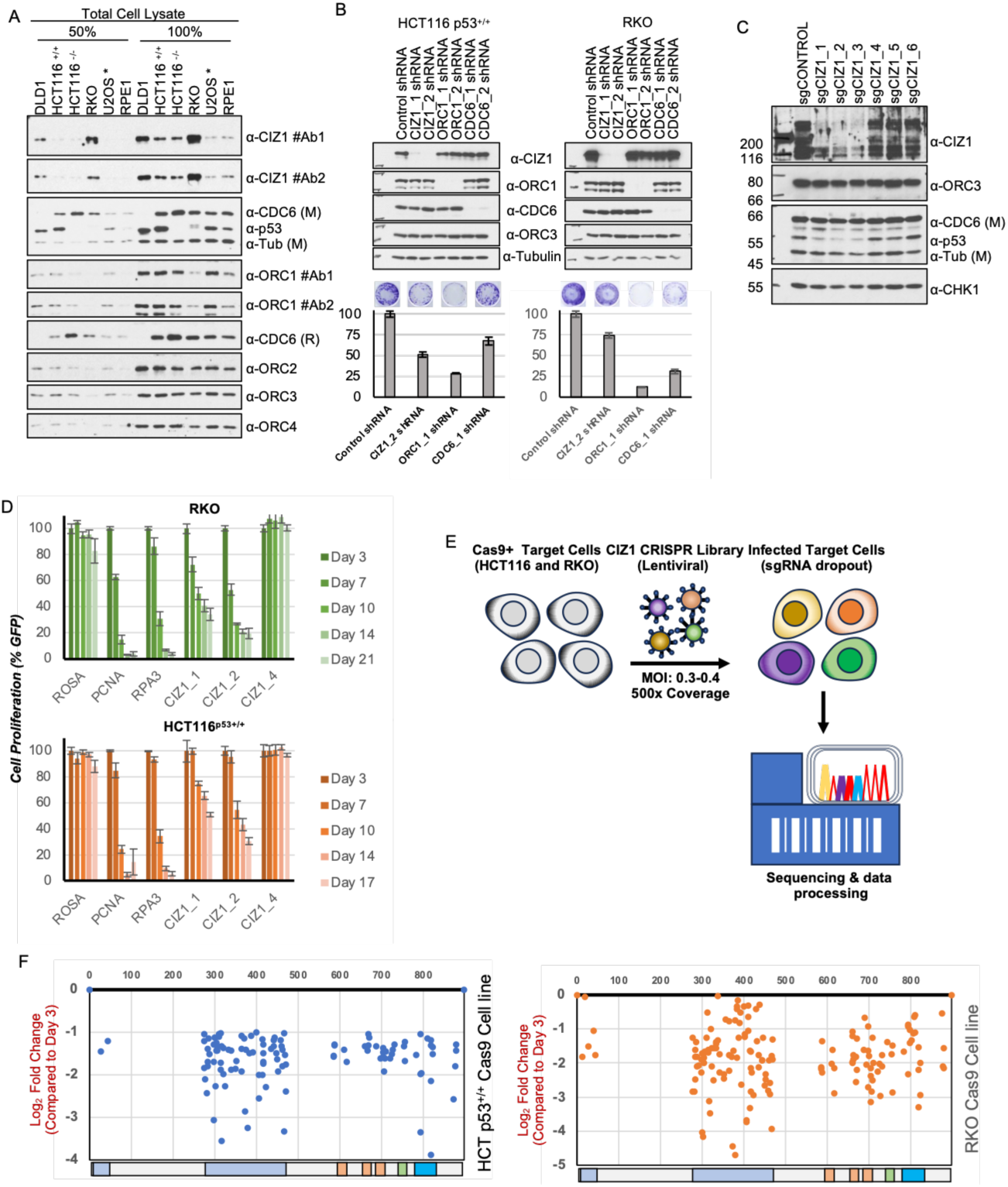
CIZ1 is required for proliferation and viability of colorectal cancer cell lines and harbors functionally essential domains identified by CRISPR screening. (A) Immunoblot analysis of total cell lysates (50% and 100% loading) from a panel of colorectal (DLD1, HCT116-p53+/+, HCT116-p53-/-, RKO), osteosarcoma (U2OS), and non-transformed (RPE1) cell lines, using two independent CIZ1 antibodies (#Ab1, #Ab2), CDC6 (monoclonal and rabbit), p53, tubulin, ORC1 (two antibodies), ORC2, ORC3, and ORC4. CIZ1 and ORC/CDC6 pre-RC components were variably expressed across cell lines, with RKO showing notably high CIZ1 levels. (B) Immunoblots (top) and clonogenic survival assays (bottom) in HCT116-p53+/+ and RKO cells expressing control, CIZ1 (two independent shRNAs), ORC1, or CDC6 shRNA. Knockdown efficiency was confirmed by immunoblotting for CIZ1, ORC1, CDC6, ORC3, and tubulin. Representative crystal violet-stained colonies and quantified colony-formation efficiency (normalized to control, mean ± SEM) show that CIZ1 depletion, like ORC1 or CDC6 depletion, significantly reduced clonogenic survival in both cell lines. (C) Immunoblot of RKO cells expressing non-targeting sgRNA (sgCONTROL) or six independent CIZ1-targeting sgRNAs (sgCIZ1_1–6), probed for CIZ1, ORC3, CDC6, p53, tubulin, and CHK1, confirming variable CRISPR-mediated CIZ1 depletion across guides. (D) Competitive proliferation assays in RKO (top) and HCT116^p53+/+ (bottom) cells expressing GFP-linked sgRNAs targeting ROSA (control), PCNA, RPA3, or CIZ1 (three independent guides: CIZ1_1, CIZ1_2, CIZ1_4), tracked as percentage GFP-positive cells over time (Days 3– 21). CIZ1_1 and CIZ1_2 guides depleted proliferating cells similarly to essential-gene controls (PCNA, RPA3), whereas CIZ1_4 behaved like the non-targeting control, indicating guide-dependent phenotypic effects. (E) Schematic of the domain-focused CRISPR knockout screening pipeline. Cas9-expressing HCT116 and RKO target cells were transduced with a CIZ1 sgRNA lentiviral library (MOI 0.3–0.4, 500x coverage), followed by sgRNA dropout selection, sequencing, and data processing to identify essential CIZ1 domains. (F) sgRNA dropout scores (log2 fold change relative to Day 3) plotted against amino acid position along CIZ1 in HCT116-p53+/+ (left, blue) and RKO (right, orange) Cas9 cell lines. Guides targeting specific domains (indicated by colored bars below each plot) showed pronounced depletion, identifying regions of CIZ1 required for cell fitness.

These results demonstrate that CIZ1 knockdown impairs DNA replication progression, leading to G2/M accumulation. Quantification of cell cycle distribution in doxycycline-induced HCT116 cells confirmed that CIZ1 knockdown increased the G1 and G2 fraction, while substantially reducing the S-phase population compared to control shRNA, collectively corroborating that CIZ1 depletion impairs S-phase entry and progression through DNA replication (Fig. 5F). Taken together, these results demonstrate that CIZ1 engages in dynamic, cell cycle regulated interactions with ORC1 and CDC6, correlating with distinct phases of replication licensing and S-phase entry. Collectively, these findings establish CIZ1 as a critical coordinator of replication machinery assembly and cell cycle progression in human cells.

### Dependence of cancer cell proliferation on CIZ1 expression levels

To characterize CIZ1 and ORC expression across cancer cell lines, total cell lysates from DLD1, HCT116 p53+/+, HCT116 p53-/-, RKO, LS174T and SW480 colorectal cancer (CRC) cells, U2OS osteosarcoma cells, and RPE1 normal diploid cells were immunoblotted at 50% and 100% loading (Fig. 6A and Fig, S6A). Two independent antibodies confirmed marked CIZ1 overexpression in RKO cells, as previously observed (Yin et al. 2013), with moderate elevation in DLD1 cells, relative to normal RPE1 and U2OS cells. Conversely, ORC1 protein levels were notably reduced in the RKO colorectal cancer line compared to the other cells. ORC2, ORC3, and ORC4 subunits were also visibly diminished (see 50% loading, Fig. 6A and Fig. S6A), suggesting a broader downregulation of the ORC complex in RKO colorectal cancer cells, potentially reflecting a compensatory response to CIZ1 overexpression. To extend the findings to breast cancer, total cell lysates from a panel of triple-negative breast cancer (TNBC) cell lines (HCC38, HCC1806, HCC1937, BT20, and MDA-MB-468) were immunoblotted for CIZ1, ORC1, ORC3, CDC6, and Cyclin A, with DDX5 serving as a loading control. CIZ1 expression was markedly elevated in HCC1806, HCC1937, and MDA-MB-468 cells, while HCC38 and BT20 showed comparatively lower levels (Fig. S6B). Notably, ORC1 levels varied with CIZ1 expression across several lines, whereas ORC3 remained relatively uniform. CDC6 and Cyclin A also exhibited variable expression (Fig. S6B). In both U2OS and RPE1 cell lines, the endogenous level of CIZ1 was very low compared to the majority of CRC and breast cancer cell lines.

To examine the consequences of over-expression of CIZ1, mGFP-CIZ1 was expressed from a doxycycline inducible promoter in U2OS cells and live-cell time-lapse imaging over approximately 22 hours revealed that CIZ1 overexpression is cyto-cytotoxic. Initially, mGFP-CIZ1 localized in discrete nuclear foci; over time, cells exhibited progressive nuclear deformation, loss of focal organization, membrane blebbing, and cell death, confirming that CIZ1 overexpression is incompatible with normal cell viability (Supplemental Movie 1). Thus, cancer cells that overexpress CIZ1 must have adapted to the higher levels.

Clonogenic survival assays were performed in HCT116 p53+/+ and RKO cells following doxycycline-induced shRNA knockdown, confirmed by immunoblotting. In HCT116 p53^+/+^ cells, CIZ1, ORC1, and CDC6 knockdown reduced colony formation to approximately 50%, 28%, and 65% of control, respectively (Fig. 6B). Strikingly, in RKO cells, which overexpress CIZ1, ORC1 knockdown caused a more dramatic reduction (to ∼12%), while CIZ1 and CDC6 knockdown reduced colonies to ∼72% and ∼30%, respectively (Fig. 6B). These findings suggest that RKO cells exhibit heightened dependency on ORC1, and that CIZ1 overexpression may confer differential sensitivity to replication factor depletion.

A series of gRNAs was developed that targeted ORC1, CDC6 or CIZ1 by selecting gRNAs in U2OS cells (e.g. Fig. 6C and Fig. S6C). These gRNAs were used to assess the functional consequences of CIZ1, ORC1, and CDC6 depletion on cancer cell proliferation. To evaluate the long-term proliferative dependency on CIZ1 in colorectal cancer cells, competitive proliferation assays were performed in RKO and HCT116 p53^+/+^ cells using CRISPR-Cas9-mediated knockout with three independent CIZ1 sgRNAs (CIZ1_1, CIZ1_2, and CIZ1_4), alongside ROSA (negative control), PCNA and RPA3 (both essential replication factors as positive control), monitored as percentage GFP over 21 days (RKO) and 17 days (HCT116 p53^+/+^) (Fig. 6D). In RKO cells, which overexpress CIZ1, sgCIZ1_1 and sgCIZ1_2 caused progressive and substantial reductions in cell proliferation, declining to approximately 35% and 20% of control by Day 21, respectively, comparable in magnitude to the essential replication factor RPA3. Notably, sgCIZ1_4 had no significant effect, consistent with it representing an inefficient guide. PCNA knockout, as expected, caused near-complete proliferative loss (Fig. 6D, top panel).

In HCT116 p53^+/+^ cells, CIZ1 knockout with sgCIZ1_1 and sgCIZ1_2 also reduced proliferation progressively, though the effect was comparatively more modest, reaching approximately 50% and 30% of control by the final timepoint, suggesting a somewhat reduced dependency relative to RKO cells. Together, these data demonstrate that CIZ1 is required for cancer cell proliferation, with RKO cells exhibiting a heightened dependency consistent with their CIZ1 overexpression status. In contrast, CIZ1^-/-^ mouse NIH3T3 cells were viable, probably due to selection of compensatory alterations in the selected cells since acute depletion in NIH3T3 cells inhibited proliferation (Coverley et al. 2005; Tollitt et al. 2026).

To map the functional domains of CIZ1 required for cancer cell proliferation, a tiling sgRNA library screen was performed across the CIZ1 coding sequence in both RKO and HCT116 cell lines. Cas9-expressing target cells were transduced with a combined CIZ1 CRISPR tiling library (463 sgRNAs) and control library (1,644 sgRNAs) at MOI 0.3–0.4, and sgRNA representation was monitored by NextSeq sequencing at multiple timepoints relative to the initial Day 3 passage, with depletion expressed as log₂ fold change (Fig. 6E). The results indicated two broad functional regions of CIZ1 in both HCT116 p53+/+ and RKO cells, with the PLD1 polyQ-rich region and the structured C-terminal domains (see Fig. 2A) being essential (Fig. 6F).

### Acute CIZ1 Depletion Disrupts Genome Organization, CTCF Occupancy, and Motility-Related Transcriptional Programs

Because CIZ1 has been implicated in nuclear matrix binding and potentially nuclear organization, we analyzed the localization of the chromosome domain organizing protein CTCF following CIZ1 depletion. A degron-tagged CIZ1 line (FKBP12^F36V^-CIZ1) was constructed that enabled rapid, dTAG-induced degradation of the endogenous protein in U2OS cells. Immunoblotting confirmed efficient loss of tagged CIZ1 within 2–6 hours of dTAG treatment, with minimal residual protein detected by anti-CIZ1 antibodies (Fig. S7), validating this system for interrogating acute, rather than chronic, consequences of CIZ1 loss. CTCF CUT&RUN before and after 12 hours of dTAG treatment was performed and CTCF occupancy was compared to a published CTCF ChIP-Seq dataset from RING1B U2OS cells and RING1B-depleted U2OS cells (Hanafiah et al. 2026). RING1b is a core enzymatic component of the Polycomb Repressive Complex 1 (PRC1). We observed CTCF binding sites at loci previously determined by ChIP-Seq, although the CUT&RUN method yielded cleaner results (Fig. 7A). At representative loci on chromosomes 1 and 11, acute CIZ1 degradation altered CTCF peak intensity, a pattern distinct from that observed upon RING1B knockdown, suggesting CIZ1 supports CTCF occupancy at specific genomic regions through a mechanism separable from Polycomb-mediated regulation (Fig. 7A). Consistent with a direct physical interaction, co-immunoprecipitation using a panel of internal CIZ1 deletion mutants mapped CTCF binding in CIZ1 to a region spanning residues 656–730, as some deletions within this interval selectively reduced co-precipitation of endogenous CTCF (Fig. 7B). These results suggest that the structured C-terminal domain of CIZ1 binds CTCF in addition to ORC1 and CDC6, Transcriptomic profiling of CIZ1-depleted cells identified both up-and downregulated genes, including CTCFL (a germline-specific paralog of the multifunctional factor CTCF) and XIST (a long coding RNA involved in X chromosome inactivation) among the most affected transcripts (Fig. 7C). Gene Ontology analysis of these differentially expressed genes revealed strong enrichment for pathways governing cell adhesion, migration, and motility (Fig. 7D), linking CIZ1 to regulation of genes governing cellular movement.

**Fig. 7.**
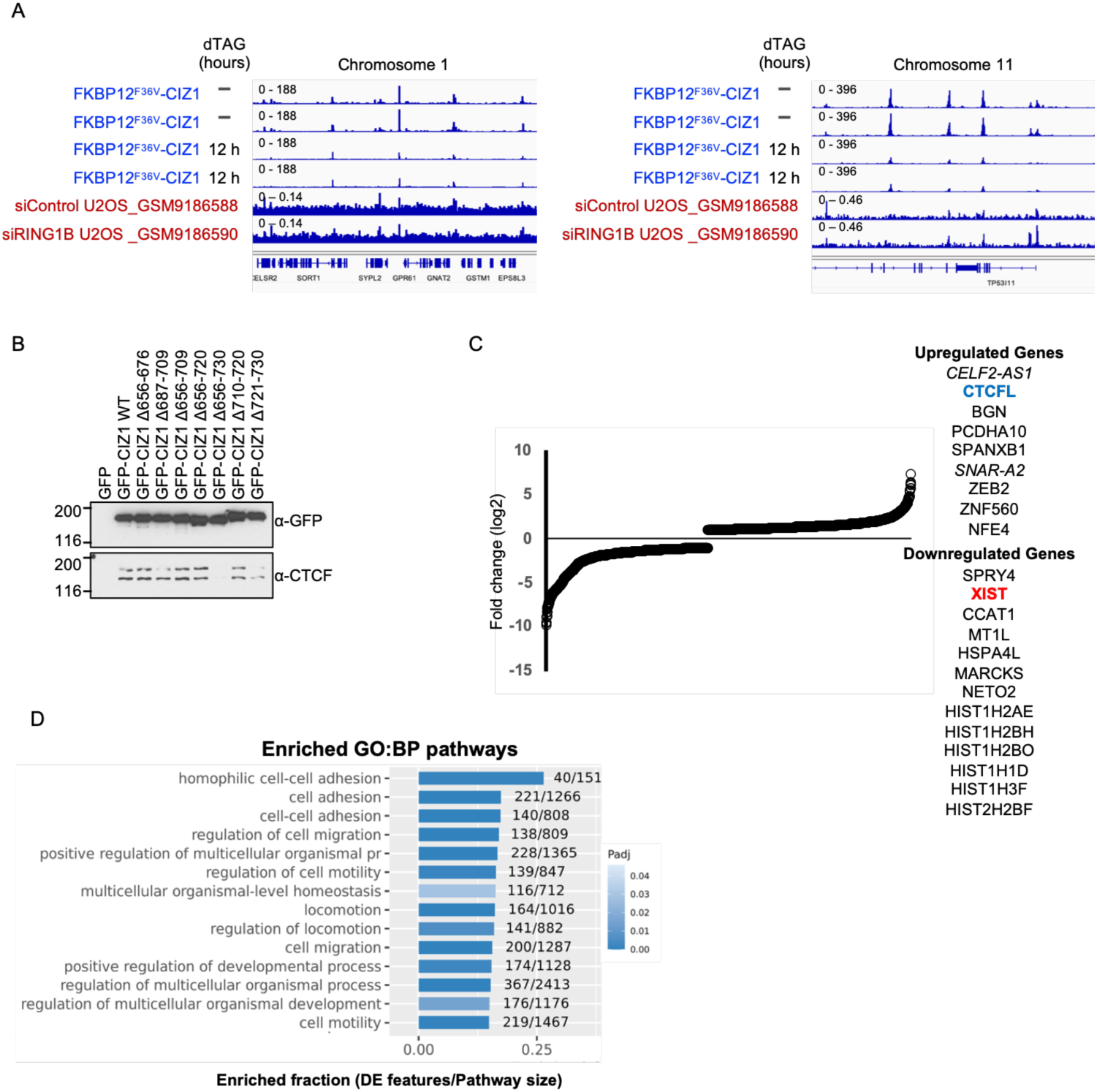
Acute CIZ1 Depletion Reveals Defects CTCF Occupancy and Motility-Related Gene Expression. (A) CTCF ChIP-seq tracks at representative loci on chromosome 1 and chromosome 11 comparing FKBP12^F36V^-CIZ1 cells before and after 12 h dTAG-induced degradation, alongside published siControl and siRING1B U2OS datasets (GSM9186588, GSM9186590). Acute CIZ1 loss produced peak changes at select CTCF-bound regions, distinct from the broader redistribution seen with RING1B knockdown, suggesting CIZ1 contributes to maintenance of a subset of CTCF binding sites independent of Polycomb-mediated mechanisms. (B) Immunoblots of GFP- and CTCF-in anti-GFP-immunoprecipitated lysates from cells expressing GFP alone, GFP-CIZ1 WT, or a panel of internal deletion mutants (Δ656–676 through Δ721–730). (C) Waterfall plot of differentially expressed genes (log2 fold change) following CIZ1 depletion, highlighting representative up- (e.g., CTCFL, ZEB2) and downregulated (e.g., XIST, histone cluster genes) transcripts. (D) Gene Ontology (Biological Process) enrichment analysis of differentially expressed genes, ranked by enriched fraction (DE features/pathway size) and colored by adjusted p-value, showing strong enrichment for cell adhesion, migration, motility, and locomotion pathways.

### CIZ1 and CDC6 are required for colorectal cancer cell proliferation and tumor growth *in vivo*

To investigate the role of CIZ1 and CDC6 in tumor growth in vivo, we first generated doxycycline-inducible shRNA stable HCT116 cell lines co-expressing firefly luciferase, enabling non-invasive bioluminescence monitoring of tumor growth following xenotransplantation into nude mice. Immunoblot validation confirmed efficient and specific knockdown of CIZ1 by CIZ1_1 and CIZ1_2 shRNAs, and CDC6 by CDC6_1 and CDC6_2 shRNAs, with tubulin serving as a loading control. Luciferase expression was confirmed across all shRNA conditions at comparable levels, validating uniform reporter expression independent of the gene knockdown (Fig. 8A). To verify that CIZ1 and CDC6 depletion impairs proliferation in this luciferase-expressing background, clonogenic survival assays were performed. CIZ1_2 shRNA and CDC6_2 shRNA both substantially reduced colony number and size compared to control shRNA cells, confirming that the proliferative defect previously observed is retained in the luciferase-expressing derivative cell line and is not an artefact of reporter integration (Fig. 8B). To further quantify the proliferative capacity of these cells, luciferase-based bioluminescence assays were performed as a surrogate for cell viability. Control shRNA cells exhibited the highest luminescence signal (∼13,500 Relative4 Light Units, RLU), while CIZ1_2 shRNA cells showed an approximately 70% reduction (∼4,000 RLU) and CDC6_2 shRNA cells showed an approximately 55% reduction (∼6,000 RLU) relative to control (Fig. 8C). The greater reduction observed upon CIZ1 knockdown compared to CDC6 knockdown suggests a potentially stronger dependency on CIZ1 for HCT116 cell proliferation.

**Fig. 8.**
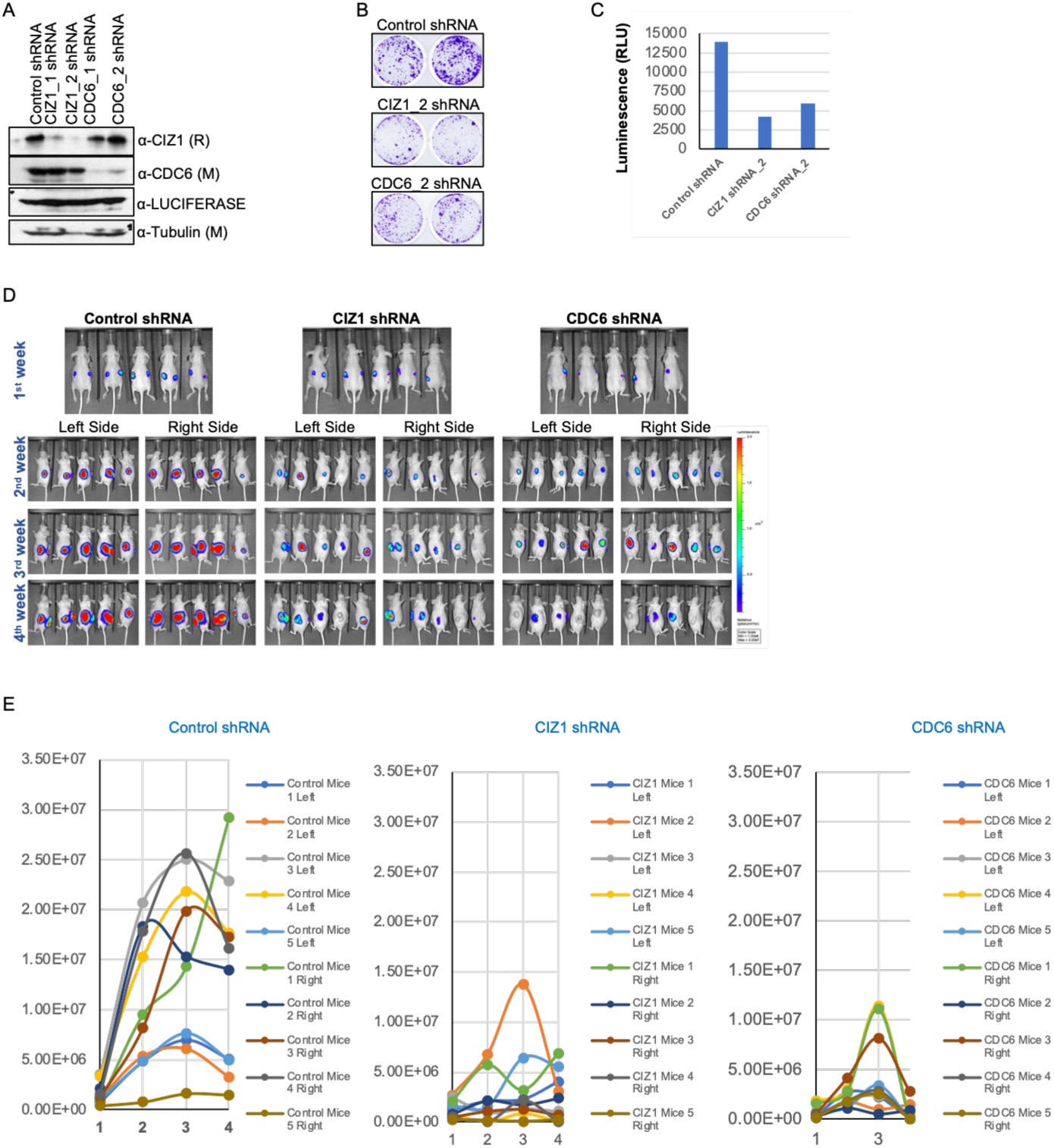
CIZ1 depletion suppresses tumor growth in vivo in a xenograft model. **N**(A) Immunoblot validation of luciferase-labeled cells expressing control, CIZ1 (two independent shRNAs, CIZ1_1 and CIZ1_2), or CDC6 (two independent shRNAs, CDC6_1 and CDC6_2) shRNA, probed for CIZ1, CDC6, luciferase, and tubulin, confirming efficient and specific knockdown prior to xenograft injection. (B) Representative crystal violet-stained colony formation assays comparing control shRNA, CIZ1_2 shRNA, and CDC6_2 shRNA cells, showing reduced colony formation upon CIZ1 or CDC6 depletion. (C) Quantification of luciferase activity (luminescence, RLU) in control shRNA, CIZ1 shRNA_2, and CDC6 shRNA_2 cells, confirming reduced luciferase signal corresponding to decreased viable/proliferating cell number following knockdown, consistent with the colony formation results in (B). (D) In vivo bioluminescence imaging of mice bearing bilateral subcutaneous xenografts of control, CIZ1, or CDC6 shRNA-expressing luciferase-labeled cells, tracked weekly from tumor implantation (1st week) through the 4th week. Left- and right-flank tumors are shown separately from week 2 onward. Control shRNA tumors exhibited progressive, robust luminescence signal expansion over time, whereas CIZ1 and CDC6 shRNA tumors showed markedly reduced and often stalled or regressing signal. (E) Quantification of tumor luminescence signal (photons/sec) over time (weeks 1–4) for individual left- and right-flank tumors in control shRNA (left), CIZ1 shRNA (middle), and CDC6 shRNA (right) cohorts (n = 5 mice/group, 10 tumors/group). Control shRNA tumors showed consistent growth to high luminescence values across nearly all tumors, while CIZ1 and CDC6 knockdown tumors displayed substantially attenuated growth kinetics and lower peak luminescence, demonstrating that CIZ1, like CDC6, is required for tumor growth in vivo.

Collectively, these data validate the luciferase-expressing doxycycline-inducible shRNA HCT116 cell lines as functionally competent tools for assessing CIZ1- and CDC6-dependent tumor growth in vivo, and establish the basis for subsequent xenograft experiments in nude mice where tumor burden can be tracked longitudinally by bioluminescence imaging.

To evaluate the requirement of CIZ1 and CDC6 for tumor growth in vivo, luciferase-expressing doxycycline-inducible shRNA HCT116 cells were subcutaneously injected bilaterally into nude mice, and tumor progression was monitored weekly by bioluminescence imaging over four weeks. In control shRNA mice, bioluminescent signal increased robustly and progressively from week 1 through week 3, with tumors on both flanks displaying intense red/yellow radiance signals indicative of vigorous tumor growth, reaching peak radiance values of approximately 2.5–3.0 × 10⁷ p/sec/cm²/sr by week 3, before a modest decline at week 4, likely reflecting tumor necrosis at advanced size (Fig. 8D and 8E). All five control mice developed bilateral tumors with consistently high signal intensity.

In striking contrast, CIZ1 shRNA mice showed markedly attenuated tumour growth throughout the experiment. Bioluminescence signals remained substantially lower across all timepoints, with the majority of tumors displaying only faint blue signals. One outlier animal (CIZ1 Mice 2 Right) showed a transient signal peak at week 3 (∼1.3 × 10⁷), which subsequently declined sharply by week 4, suggesting incomplete knockdown or escape in that individual. Overall, CIZ1 depletion resulted in a dramatic suppression of tumor establishment and growth compared to controls.

Similarly, CDC6 shRNA mice exhibited significantly reduced bioluminescence signals relative to controls, with peak radiance values generally not exceeding 1.0 × 10⁷ p/sec/cm²/sr and several tumors remaining near background levels throughout. The inhibitory effect of CDC6 knockdown, while significant, appeared somewhat less uniform than CIZ1 knockdown, with greater inter-animal variability observed.

Collectively, these in vivo data demonstrate that both CIZ1 and CDC6 are required for efficient HCT116 tumor growth in nude mice, strongly supporting a functionally important role for CIZ1, alongside CDC6, in sustaining tumor proliferation in a colorectal cancer xenograft model.

## Discussion

CIZ1 has been implicated in tumorigenesis and consistent with previous results (Pauzaite et al. 2016) we found variable expression of CIZ1 in various colon and breast cancer cells and essential for maintenance of tumor growth in vivo. The levels of CIZ1 varied with the levels of ORC1, but not ORC3 and ORC4, suggesting mutual regulation.

The interaction of human CIZ1 protein with ORC1, CDC6 and Cyclin-CDK2 and Cyclin A-CDK2 and its requirement for efficient pre-RC assembly is consistent with a role in DNA replication in a variety of human cells. Previous studies using isolated mouse cell nuclei have implicated the N-terminus as of CIZ1 as stimulating DNA replication (Ainscough et al. 2007; Copeland et al. 2010; Coverley et al. 2005; Copeland et al. 2015; YIN et al. 2013; Wilson and Coverley 2013). In contrast, we find that both the N-terminal and C-terminal domains of human CIZ1 are required for cell proliferation. An interaction between CDC6 and the N-terminus of CIZ1 has been demonstrated in mouse cells and Cyclin A-CDK2 phosphorylation did not disrupt the interaction (Copeland et al. 2015; Coverley et al. 2005). In direct contrast, we observed that the C-terminal Zn-finger region of CIZ1 binds to both ORC1 and CDC6 and Cyclin A-CDK2 disrupts both interactions. One possibility IDR in the N-terminus of CIZ1 and the IDR in CDC6 interact under certain biochemical conditions used previously, but the mutant data we obtained suggest that it is the C-terminal domain of CIZ1 that mediates ORC1 and CDC6 interactions.

The interaction between CIZ1 and ORC1 and CDC6 is regulated by CDKs. We find that Cyclin A-CDK2, but not Cyclin E-CDK2 kinase activity blocks the interaction between CIz1 and ORC1, whereas both CDKs block the interaction between CIZ1 and CDC6. The latter result is in contract to the conclusion that Cyclin A-CDK2 does not block the interaction between CIZ1 and CDC6 (Copeland et al. 2015), however, the conclusion was based on phospho-mimetic mutants of CIZ1 whereas we showed direct inhibition by kinase activity.

Notably, however, CIZ1 associates with ORC1 via its N-terminal intrinsically disordered region containing the Cy motif (ORC1²³⁵KRL²³⁷), a site that is spatially and functionally distinct from the canonical chromatin-binding BAH domain of ORC1 (Frisbie and Bleichert 2026; Stillman et al. 2025; Parker et al. 2019). The fact that Cyclin E-CDK2 does not prevent binding of ORC1 to CIZ1 would allow the CDC6-Cycin E-CDK2 complex to bind to ORC1 and CIZ1 for pre-RC assembly. Upon activation of Cyclin E-CDK2 kinase activity, CDC6 could be released allowing formation of the double hexamer of MCM and the pre-RC. The observation that both ORC1 and CDC6 binding to CIZ1 is absent in G2/M phase suggests that Cyclin A-CDK2 activity in late G2/M phase prevents these interactions with CIZ1, just like ORC1-CDC6 interaction in mitosis is blocked by Cyclin A-CDK2 (Hossain et al. 2021). Such activity might also release of CIZ1 from mitotic chromosomes.

CDC6 interacts with CIZ1 through its conserved Cy motif (CDC6⁹⁴RRL⁹⁶), previously shown to mediate binding to G1/S Cyclin-CDK and ORC1 (Borlado and Méndez 2008; Petersen et al. 2000; Hossain et al. 2021; Jiang et al. 1999). Importantly, mutations within the CDC6 Cy motif that disrupt interactions with Cyclin A–CDK2 or Cyclin E–CDK2 also abolish CIZ1 binding, revealing an interconnected regulatory network in which CIZ1, Cyclin-CDK complexes, and CDC6 may compete for or coordinately engage a shared interaction surface. This finding is mechanistically significant because Cy motif-dependent binding of CDC6 to Cyclin–CDK complexes is known to promote CDC6 phosphorylation and its subsequent nuclear export during S phase (Petersen et al. 2000; Jiang et al. 1999). Therefore, CIZ1 most likely functions as a spatial regulator of replication licensing by modulating CDC6 accessibility to ORC1 and Cyclin-CDK complexes and controlling its retention or release at the nuclear matrix in a cell cycle-dependent manner.

Consistent with the dynamic exchange of ORC1 and CDC6 with matrix-bound CIZ1, In vitro competition assays demonstrated that increasing concentrations of ORC1 progressively disrupted the CIZ1-CDC6 interaction, whereas the reciprocal inhibitory effect of CDC6 on CIZ1-ORC1 binding was comparatively modest. This asymmetric competition suggests a hierarchical mechanism governing replication complex assembly, in which ORC1 preferentially associates with CIZ1 during pre-replication complex (pre-RC) formation in early G1 phase, while CDC6 binding becomes more prominent during late G1 phase when additional pre-RC assembly occurs. Such a model is consistent with the established temporal order of pre-RC assembly, whereby ORC recruits CDC6, which subsequently promotes loading of the MCM2–7 helicase complex (Stillman et al. 2025; Costa and Diffley 2022; Frisbie and Bleichert 2026). Within this framework, CIZ1 may function as a dynamic and competitive scaffold that facilitates the ordered exchange between ORC1- and CDC6-containing complexes, facilitated by the sequential activity of Cyclin E-CDK2 and then Cyclin A-CDK2. This proposed role places CIZ1 at a critical regulatory node linking replication licensing and activation, while also introducing an additional level of spatial organization on the nuclear matrix (Ainscough et al. 2007).

Beyond its established role in replication licensing, we observed that CIZ was required for maintaining CTCF on chromatin at discrete binding sites. This interpretation is reinforced by the mapping of a direct CIZ1-CTCF interaction to a discrete internal region (residues 720–730), a domain overlapping the C-terminal ZnF2/ZnF3-Cy motif interface that mediates CIZ1 engagement with ORC1 and CDC6. CIZ1 may therefore serve as a multivalent nuclear matrix scaffold with functionally segregated interfaces, coordinating replication licensing through its C-terminal domain while supporting CTCF occupancy, and by extension chromatin loop or topologically associating domain boundaries, through this more internal region.

The transcriptional consequences of acute CIZ1 depletion further support a functional, rather than incidental, link between CIZ1 and CTCF-dependent chromatin architecture. Among the most affected transcripts were CTCFL, the testis-specific paralogue of CTCF implicated in aberrant chromatin remodeling when mis-expressed in somatic tissue, and XIST, a locus whose regulation is itself dependent on CTCF-anchored chromatin loops, consistent with previous reports linking CIZ1 to X chromosome inactivation maintenance (Sofi and Coverley 2023; Ridings-Figueroa et al. 2017; Dobbs et al. 2023). More broadly, differentially expressed genes were strongly enriched for pathways governing cell adhesion, migration, and motility, phenotypes not classically associated with replication licensing factors. The CTCF-linked transcriptional program raises the possibility that CIZ1 contributes to malignant progression through a second, architecturally mediated axis, one that shapes the invasive or migratory potential of cancer cells independently of its role in sustaining replication. Distinguishing whether these motility-associated transcriptional changes arise directly from altered CTCF occupancy at their regulatory loci, or indirectly through broader chromatin or replication stress-associated effects of CIZ1 loss, will be an important focus of future work, as will determining whether the CIZ1-CTCF interaction is constitutive or itself regulated across the cell cycle in a manner analogous to the ORC1 and CDC6 interactions described above.

## Acknowledgements

The research was supported by grants to BS from the National Institute of General Medical Sciences (GM45466) and the National Cancer Institute (CA13106), as well as the Cold Spring Harbor Laboratory Cancer Center Grant (CA45508). Research in the Stillman Laboratory is supported by the Goldring Family Foundation. We thank Dr. Osama El Demerdash for computational support and Matthew Tuttle for technical support.

## Materials and Methods

### Cell culture

HCT116 (p53+/+), U2OS, RPE-1, HeLa and HEK293/HEK293T cell lines were cultured in DMEM (Gibco) supplemented with 10% fetal bovine serum and 1% penicillin/streptomycin at 37°C in a humidified incubator with 5% CO2. RKO, DLD1, and the triple-negative breast cancer lines HCC38, HCC1806, HCC1937, BT20, and MDA-MB-468 were cultured in RPMI-1640 or DMEM as appropriate for each line, supplemented with 10% fetal bovine serum and 1% penicillin/streptomycin. HeLa spinner suspension cells used for cell-cycle synchronization were maintained in Joklik’s modified MEM supplemented with 5% newborn calf serum.

Tetracycline-inducible U2OS cell lines stably expressing mGFP-CIZ1 or mGFP-ORC1 (Flp-In T-REx system) were maintained in 10% Tet-system-approved fetal bovine serum (Takara/Clontech) and appropriate selection antibiotics (hygromycin B, blasticidin), and induced with 1 µg/mL tetracycline or doxycycline as indicated. Doxycycline-inducible shRNA HCT116 (p53+/+) and RKO cell lines were maintained in 10% Tet-system-approved fetal bovine serum and induced with 0.5–1 µg/mL doxycycline for the times indicated in the figure legends. All cell lines were routinely tested for mycoplasma contamination and confirmed negative.

### Plasmid construction and mutagenesis

Full-length human CIZ1 and full-length mouse Ciz1 cDNAs, together with a series of N- and C-terminal truncations (human: 1–550, 550–898, 1–835, 1–782, 1–740, 1–686, 1–655, 1–627, 1–594, 1–527, 1–435, 594–740, 665–740, 675–740, 685–740, 695–740, 655–730, 655–720, 655–710, 655–685; mouse: 1–550, 550–845), internal deletions (Δ656–676, Δ656–685, Δ656–709, Δ656–710, Δ656–720, Δ656– 730, Δ721–730), point mutants (LxF A-A; Cy1 ^714^KEL^716^ A-A; Cy2 ^717^KSL^719^ A-A; Cy3 ^820^KSL^822^ A-A; single, double, and triple Cy-motif combinations, Cy12 and Cy123; CDK phospho-site mutants T585A, T696A, T876A and double/triple combinations, designated TA12/TA13/TA23/TA123), and zinc-finger deletions (ΔZn2, ΔZn3, ΔZn2+3) were generated by PCR-based cloning into pGEX-6P1 (GST fusion, bacterial expression), pMAL-c2E (MBP fusion, bacterial expression), pEGFP-C1 (GFP fusion, mammalian expression), and pmCherry-C1 (mCherry fusion, mammalian expression) vectors. A T7 and HA epitope tag was appended to selected MBP-CIZ1 constructs (T7-MBP-CIZ1 or HA-MBP-CIZ1) for competition and reconstitution assays.

Recombinant human ORC1 and its mutants (M5A: ^232^RARKR^236^→^232^AAAAA^236^; A-A: ^235^ARA^237^; BP4; BP5; Cy-motif mutant ORC1-Cy), together with ORC2, ORC3, ORC4, and CDC6 and its mutants (Cy-motif AAL: ^94^RRL^96^→^94^AAL^96^; K92A, R94A, R95A, L96A and combinations) were cloned into pMAL-c2E for MBP-fusion bacterial expression. A series of N-terminal ORC1 truncations (1–100, 1–200, 1–300, 1– 400, ΔN93, ΔN200, ΔN300, ΔN400, 1–861 full length, 300–861) and an ORC1 internal fragment (100– 250, harboring point mutants TP230/1AA, R232A, R234A, K235A, R236A, L237A, KL235/7AA, EL238/9AA) were similarly generated in pMAL-c2E. Human CDC6 and its mutants were also cloned into pGEX-6P1 to produce GST-fusion proteins. Human ORC1 WT and mutants were additionally cloned into pEGFP-C1 for transient overexpression in mammalian cells. Human and mouse CDK inhibitors p21^Cip1/Waf1^ and p27^Kip1^ were cloned into pMAL-c2E as MBP fusions.

For the acute-degron system, the endogenous CIZ1 locus was tagged at the N-terminus with an HA-FKBP12F36V degron cassette by CRISPR-Cas9-mediated homology-directed knock-in, generating an FKBP12F36V-CIZ1 fusion allele compatible with dTAG-mediated degradation.

All point mutations, internal deletions, and truncations were generated by site-directed mutagenesis using Phusion high-fidelity DNA polymerase (NEB). Internal deletions were generated using phosphorothioate-modified PCR primers followed by T7 gene 6 exonuclease digestion of the unmodified parental strand, as previously described ((Stoynova et al. 2004). All constructs were verified by Sanger sequencing across the full insert. Oligonucleotide sequences used for cloning and mutagenesis are available upon request.

### Antibodies

The following primary antibodies were used for immunoblotting, immunoprecipitation, and immunofluorescence: rabbit polyclonal anti-CIZ1 [two independent antibodies, designated #Ab1 (PCRP-CIZ1-1A2, DSHB) and #Ab2 (PA5-41533, Thermo)]; rabbit polyclonal anti-CTCF (D31H2, Cell Signaling); mouse monoclonal anti-GFP (A11120; Invitrogen); rabbit polyclonal anti-CDC6 (CS1881, CSHL facility) and mouse monoclonal anti-CDC6; two independent anti-ORC1 antibodies (#1 and #2, including 78-1-172; Kara et al., 2015); rabbit polyclonal anti-ORC2; rabbit polyclonal anti-ORC3; rabbit polyclonal anti-ORC4; mouse monoclonal anti-MCM2 (#610700; BD Biosciences); rabbit polyclonal anti-Cyclin A (A305-254A; Bethyl); rabbit polyclonal anti-Cyclin E; mouse monoclonal anti-p53 (DO-1); mouse monoclonal anti-α-tubulin; rabbit polyclonal anti-CHK1; rabbit polyclonal anti-DDX5; mouse monoclonal anti-HA; mouse monoclonal anti-T7 (CSHL facility); mouse monoclonal anti-MBP; mouse monoclonal anti-GST; and mouse monoclonal anti-luciferase. Species-appropriate HRP-conjugated secondary antibodies (Jackson ImmunoResearch or Cell Signaling Technology) were used for immunoblot detection, and Alexa Fluor-conjugated secondary antibodies (Invitrogen) were used for immunofluorescence and flow cytometry.

### Expression and purification of recombinant proteins

MBP- and GST-fusion proteins were expressed and purified from E. coli BL21 using amylose resin or glutathione-Sepharose beads, respectively, following a previously described protocol (Hossain and Stillman 2012). Briefly, transformed bacterial cultures were grown in LB medium at 37°C to an OD600 of 0.7–0.9 and induced with 0.3 mM IPTG for 12 h at 16°C. Cell pellets were lysed by sonication in Lysis Buffer A (25 mM Tris-HCl pH 7.5, 150 mM NaCl, 0.02% NP-40, 5 mM benzamidine-HCl, 1 mM PMSF, cOmplete protease inhibitor cocktail [Roche], 10% glycerol) supplemented with 100 µg/mL lysozyme. Clarified lysates were incubated with pre-equilibrated amylose or glutathione agarose beads for 3 h at 4°C. Bead-bound protein was washed with ten column volumes of Buffer A containing 500 mM NaCl and 0.05% NP-40, followed by ten column volumes of Buffer A alone. Proteins were eluted stepwise with Buffer A containing 20 mM maltose (MBP fusions) or 20 mM reduced glutathione, pH 7.5 (GST fusions). Peak fractions were pooled, concentrated, and dialyzed against storage buffer, and protein concentration was determined by Bradford assay. Purified protein purity and integrity were verified by Coomassie-stained SDS-PAGE. Cyclin A–CDK2 and Cyclin E–CDK2 kinase complexes were expressed and purified from baculovirus-infected Sf9 insect cells as previously described (Hossain and Stillman 2012).

### In vitro binding (pull-down) assays

For MBP, GST, and T7 pull-down assays, anti-MBP, anti-GST, or anti-T7 antibodies were pre-bound to Protein G Dynabeads and incubated with the indicated purified MBP- or GST-fusion proteins in Binding Buffer (25 mM Tris-HCl pH 7.5, 150 mM KCl, 0.15% Nonidet P-40, 0.1 mM EDTA, 5 mM magnesium acetate, 1 mM DTT). Bait proteins immobilized on beads were incubated with the indicated prey proteins (GST-CIZ1, MBP-CIZ1, MBP-ORC1, GST-CDC6, MBP-CDC6, Cyclin E-CDK2, Cyclin A-CDK2, and their respective mutants) for 4–5 h at 4°C with end-over-end rotation. Beads were washed three times with Binding Buffer, and bound proteins were eluted in Laemmli sample buffer, resolved by SDS-PAGE, and detected by immunoblotting with the antibodies indicated in the figure legends.

For the ORC1–CDC6 competition assay, purified T7-MBP-CIZ1 (70 nM) was incubated with a fixed concentration of HA-MBP-CDC6 (70 nM) or HA-MBP-ORC1 (70 nM) in the presence of increasing concentrations (0, 70, 140, 280 nM) of the reciprocal competitor protein. Complexes were captured with anti-T7 antibody-bound beads, washed, and analyzed by immunoblotting with anti-HA antibody to quantify relative ORC1 and CDC6 recruitment.

### In vitro kinase assays

Purified GST-CIZ1 (wild-type, phospho-site mutants, or Cy-motif mutants; full-length or C-terminal fragments) was incubated with purified Cyclin E–CDK2 or Cyclin A–CDK2 kinase complexes in Kinase Buffer (50 mM Tris-HCl pH 7.5, 10 mM MgCl2, 1 mM DTT) supplemented with ATP (including [γ-32P]-ATP or cold ATP as appropriate for downstream detection) for 30 min at 30°C. Reactions were terminated with Laemmli sample buffer, resolved by SDS-PAGE, and substrate loading was confirmed by Coomassie staining of the same gel. Phosphorylation was visualized by autoradiography/phosphorimaging (Typhoon imaging system).

### Immunoprecipitation and immunoblotting

For transient expression, HEK293 or HEK293T cells were transfected with the indicated plasmids using Lipofectamine 2000 (Thermo Fisher Scientific) according to the manufacturer’s instructions. For immunoprecipitation, cells were harvested, washed in PBS, and lysed in IP Lysis Buffer (20 mM Tris-HCl pH 7.5, 300 mM NaCl, 0.3% NP-40, 5 mM MgCl2, 0.1 mM EDTA, 10% glycerol, 1 mM DTT, 1 mM CaCl2, 20 µM MG132, and protease/phosphatase inhibitor tablets [Roche]) supplemented with Benzonase (Sigma-Aldrich) and incubated for 30 min on ice. Lysates were diluted to 150 mM NaCl/0.15% NP-40 with Dilution Buffer and clarified by centrifugation at 14,000 rpm for 15 min at 4°C. GFP-tagged proteins were immunoprecipitated with anti-GFP antibody for 4 h, followed by a 2 h incubation with pre-washed Protein G Dynabeads at 4°C. Beads were washed three times in Wash Buffer (20 mM Tris-HCl pH 7.5, 150 mM NaCl, 0.15% NP-40, 5 mM MgCl2, 0.1 mM EDTA, 10% glycerol, 1 mM DTT, protease/phosphatase inhibitors), eluted in Laemmli sample buffer, and resolved on 8% SDS-PAGE gels for immunoblotting.

For endogenous co-immunoprecipitation of CIZ1-interacting proteins from whole-cell or chromatin-enriched extracts, asynchronous or synchronized U2OS or HeLa cells were lysed in Lysis Buffer A (20 mM Tris-HCl pH 7.5, 250 mM NaCl, 5 mM MgCl2, 0.1 mM EDTA, 0.3% NP-40, 1 mM DTT, 1 mM CaCl2, 5% glycerol, supplemented with Benzonase, roscovitine, MG132, and protease/phosphatase inhibitor cocktails) for 30 min at 4°C with rotation. Extracts were diluted with an equal volume of Dilution Buffer B (Buffer A lacking NaCl and NP-40) to a final concentration of 125 mM NaCl/0.15% NP-40 and clarified by centrifugation. Clarified extracts were incubated with anti-CIZ1 (two independent antibodies), anti-CDC6, anti-ORC1, or isotype-matched IgG control antibodies pre-bound to Protein G Dynabeads. Beads were washed, and bound proteins were eluted in 2× Laemmli sample buffer for immunoblot analysis with the antibodies indicated in the figure legends.

For cell-cycle-resolved co-immunoprecipitation, HeLa spinner suspension cells were synchronized by double-thymidine block, released, and harvested at 2 h intervals from 0–24 h post-release; extracts were prepared as above and subjected to CIZ1 or IgG immunoprecipitation followed by immunoblotting for CIZ1, ORC1, CDC6, ORC3, ORC4, MCM2, and Cyclin A.

For CIZ1-CTCF co-immunoprecipitation, cells expressing GFP alone, GFP-CIZ1 WT, or the indicated internal CIZ1 deletion mutants were lysed as above, and GFP- or endogenous CTCF-immunoprecipitates were analyzed by immunoblotting with anti-GFP and anti-CTCF antibodies.

### Cell synchronization and cell-cycle analysis

For double-thymidine block and release, HeLa spinner or adherent cells were incubated with 2 mM thymidine for 18 h, released into fresh medium for 9 h, and re-blocked with 2 mM thymidine for 16 h. Following release from the second block, cells were harvested at the indicated time points (0–24 h). DNA content was analyzed by propidium iodide or FxCycle Violet (Thermo Fisher Scientific) staining and flow cytometry to confirm synchrony and to assign cell-cycle phase (G1/S, S, G2/M, G1) at each collection time.

### EdU incorporation, immunofluorescence, and co-localization

For colocalization studies, mGFP-CIZ1-expressing U2OS cells were pulse-labeled with 10 µM EdU for 60 min, fixed in 4% paraformaldehyde (PFA) in PBS for 15 min, permeabilized with 0.5% Triton X-100, and processed using the Click-iT EdU Alexa Fluor 594/647 Imaging Kit (Thermo Fisher Scientific) per the manufacturer’s protocol. Nuclei were counterstained with DAPI. Images were acquired on a PerkinElmer spinning-disk confocal microscope equipped with a Nikon Ti-E inverted stand and a 40–60× oil-immersion objective, and presented as single-plane or maximum-intensity z-projections.

For CIZ1-ORC1 co-localization, tetracycline-inducible mGFP-ORC1 stable U2OS cells were transiently transfected with mCherry-CIZ1 using Lipofectamine 2000. Cells were examined with or without prior extraction with 0.5% Triton X-100 in CSK buffer (10 mM PIPES pH 6.8, 100 mM NaCl, 300 mM sucrose, 1 mM EGTA, 1 mM MgCl2, 1 mM DTT, protease/phosphatase inhibitors) on ice for 5 min to remove soluble, non-matrix-bound protein, prior to fixation in 4% PFA and imaging at 40× magnification.

### Flow cytometry: EdU/MCM/CIZ1 cell-cycle profiling

Asynchronous or synchronized cells were pulse-labeled with 10 µM EdU for 60 min prior to harvest. For chromatin-bound protein analysis, cells were pre-extracted in CSK buffer containing 0.5% Triton X-100 with protease/phosphatase inhibitors on ice for 5 min, then fixed in 4% PFA for 15 min. EdU was detected using the Click-iT Plus EdU Alexa Fluor 488 or 647 Flow Cytometry Assay Kit (Thermo Fisher Scientific, C10635/C10420) per the manufacturer’s protocol, substituting 1% BSA-PBS + 0.1% NP-40 for the kit wash buffer where indicated. Chromatin-bound MCM2 was detected using anti-MCM2 antibody (#610700, BD Biosciences) followed by Alexa Fluor 647-conjugated donkey anti-mouse secondary antibody (Jackson ImmunoResearch); CIZ1 was detected with a fluorophore-conjugated anti-CIZ1 antibody (610_20 YG channel). DNA content was assessed with FxCycle Violet Stain (Thermo Fisher Scientific, F10347). A minimum of 10,000 cells were acquired per condition on a BD LSRFortessa Dual SORP (BD Biosciences), with GFP/Alexa Fluor 488 collected off the 488 nm laser (505LP/530-30 filter), Alexa Fluor 647 off the 638 nm laser (670/30 filter), and FxCycle Violet off the 407 nm laser (450/50 filter). Gating for negative populations was established using unstained and secondary-antibody-only controls. Data were analyzed using FlowJo software.

### Generation of doxycycline-inducible shRNA cell lines

shRNA sequences targeting CIZ1, ORC1, or CDC6 (two independent hairpins per gene) and a non-targeting control shRNA were cloned into a doxycycline-inducible lentiviral vector (Tet-pLKO-puro, AddGene # 21915). Lentivirus was produced in HEK293T cells (see Viral production and transduction below) and used to transduce HCT116 (p53+/+) and RKO cells, followed by antibiotic selection to generate stable, inducible knockdown lines. Knockdown was induced with 0.5–1 µg/mL doxycycline for 48–96 h and validated by immunoblotting for the corresponding target protein prior to functional assays.

### CRISPR/Cas9 knockout cell line generation

To generate stable CIZ1 knockout U2OS cells, parental U2OS cells were first transduced with Cas9-expressing lentivirus and selected with puromycin to generate U2OS-Cas9 cells. These cells were subsequently transduced with individual CIZ1-targeting sgRNA lentiviruses (six independent guides, sgCIZ1_1–6) or a non-targeting control sgRNA (sgCONTROL) and selected with blasticidin to generate stable, double-antibiotic-resistant knockout populations. CIZ1 depletion was confirmed by immunoblotting.

### Domain-focused CRISPR tiling library screen

#### Guide design

A tiling sgRNA library targeting the full CIZ1 coding sequence was designed by extracting every possible guide directly upstream of an NGG PAM across the CIZ1 exons. Candidate guides were aligned to the GRCh38 genome using the BatMis exact k-mismatch aligner (Tennakoon et al. 2012), allowing up to three mismatches for off-target evaluation. Off-target sites were penalized according to mismatch number and position, and an aggregate off-target score was calculated for each guide (Hsu et al. 2013). The final tiling library comprised 463 CIZ1-targeting sgRNAs, combined with a control library of 1,644 sgRNAs (non-targeting and essential/non-essential gene controls).

#### Library cloning

sgRNA oligonucleotides, including all controls, were synthesized on a pooled array platform (Twist Bioscience) and cloned into a BsmBI-digested U6-sgRNA-GFP lentiviral vector (LRG2.1-derived, Addgene #108098) by Gibson Assembly, as previously described (Tarumoto et al. 2018). Library representation and cloning fidelity were confirmed by MiSeq sequencing (Illumina), confirming that >95% of designed guides were represented within five-fold of the mean abundance.

#### Screening

Cas9-expressing HCT116 (p53+/+) and RKO target cells were transduced with the pooled CIZ1 tiling and control sgRNA libraries at an MOI of 0.3–0.4 to ensure single-copy integration, with sufficient cell numbers infected to maintain ≥500× library coverage. Cells were harvested at day 3 post-infection as the initial reference time point and cultured for the durations indicated in the figure legends (up to Day 14–21), with genomic DNA extracted using the QIAamp DNA Midi Kit (QIAGEN).

#### Sequencing and analysis

The sgRNA cassette was PCR-amplified from genomic DNA, and Illumina-compatible adapters and sample-specific barcodes were added by a second PCR. Libraries were quantified by Agilent Bioanalyzer, pooled equimolarly, and sequenced on a NextSeq 500/550 (single-end, 76 cycles). sgRNA counts were quantified by exact-match alignment to barcode and guide sequences. Guide-level and gene-level depletion (log2 fold change relative to Day 3) was calculated using MAGeCK (Li et al. 2014), which applies median normalization and negative binomial modeling. Essential domains were identified from tiling data using ProTiler (He et al. 2019), which applies denoising and wavelet-based change-point detection to define CRISPR knockout hypersensitive (CKHS) regions.

### Competitive proliferation (GFP dropout) assay

Cas9-expressing RKO and HCT116 (p53+/+) cells were transduced with individual GFP-linked sgRNA lentiviruses (ROSA negative control; PCNA and RPA3 positive controls; or CIZ1-targeting guides CIZ1_1, CIZ1_2, CIZ1_4) at an MOI of 0.3–0.4. Beginning on day 3 post-transduction (P1), cells were passaged every 2–3 days and the percentage of GFP-positive cells was measured at each passage by flow cytometry (Guava easyCyte) up to Day 17–21, in three technical replicates per timepoint across three independent biological replicates. Values were normalized to the GFP percentage at the initial (P1) timepoint for each guide.

### Clonogenic survival (colony formation) assay

Cells expressing control, CIZ1, ORC1, or CDC6 shRNA (or the corresponding sgRNA where indicated) were seeded at low density in 6-well plates following knockdown/knockout validation and cultured for 10– 14 days, with medium (± doxycycline) refreshed every 3–4 days. Colonies were fixed with methanol and stained with 0.5% crystal violet in 20% methanol. The amount of dye retained by the colonies in each well was quantified by destaining with 1 mL of 10% glacial acetic acid for 30 minutes, followed by measuring the absorbance of the destaining solution at 595 nm (OD₅₉₅) using an Eppendorf Biophotometer.

### Generation and validation of the FKBP12^F36V^-CIZ1 degron cell line and dTAG-mediated degradation

An FKBP12^F36V^-HA degron tag was knocked in at the endogenous CIZ1 N-terminus by CRISPR-Cas9-mediated homology-directed repair to generate an FKBP12F36V-CIZ1 fusion protein in the parental cell line. Clones were validated by PCR genotyping and immunoblotting with anti-HA and anti-CIZ1 antibodies, confirming a molecular weight shift consistent with the predicted tagged protein (endogenous CIZ1, ∼99 kDa; degron tag, ∼14 kDa; tagged CIZ1, ∼113 kDa). For acute CIZ1 depletion, cells were treated with 500 nM dTAG-13 (or the equivalent dTAG degrader) for the indicated durations (2, 6, or 12 h), with DMSO as vehicle control. Degradation efficiency was confirmed by immunoblotting with anti-HA and anti-CIZ1 antibodies at each time point prior to downstream ChIP-seq or RNA-seq analysis.

### CTCF CUT&RUN and data analysis

CUT&RUN was performed on FKBP12F36V-CIZ1 cells before (DMSO) and after 12 h of dTAG-induced CIZ1 degradation (two biological replicates per condition), using a protocol adapted from (Meers et al. 2019). Briefly, cells were harvested, washed, and bound to concanavalin A-coated magnetic beads. Cells were permeabilized with digitonin and incubated overnight at 4°C with anti-CTCF antibody (or IgG control) diluted in antibody buffer. Following primary antibody incubation, cells were washed and incubated with pA/G-MNase fusion protein to tether MNase to antibody-bound CTCF sites. Chromatin digestion was activated by addition of CaCl₂ and carried out at 0°C to restrict cleavage to antibody-tethered sites, after which the reaction was stopped and CTCF-bound chromatin fragments were released and diffused into solution. *E. coli* spike-in DNA was added at this stage for normalization. DNA was purified using standard phenol-chloroform extraction or column-based purification and used to construct sequencing libraries, which were sequenced on an Illumina platform. Reads were aligned to the human genome (GRCh38/hg38) using bowtie2, with spike-in reads aligned in parallel to the *E. coli* genome for normalization of sequencing depth across samples. CTCF peaks were called using SEACR with IgG as the background control.

To confirm that CTCF peaks identified in our CUT&RUN dataset represent bona fide CTCF binding events, we compared our peak calls against publicly available siControl and siRING1B CTCF ChIP-seq datasets from U2OS cells (GSM9186588, GSM9186590), downloaded from the Gene Expression Omnibus (GEO). These datasets were processed using the same alignment pipeline (bowtie2 alignment to GRCh38/hg38) and peak overlap was assessed to validate the specificity of our CTCF peak calls. Aligned tracks were visualized using the Integrative Genomics Viewer (IGV).

### RNA-seq and differential gene expression analysis

Total RNA was extracted from control (DMSO) and dTAG-treated (12 h) FKBP12F36V-CIZ1 cells using the RNeasy Mini Kit (QIAGEN), and RNA integrity was confirmed by Bioanalyzer. Poly(A)-selected sequencing libraries were prepared and sequenced on an Illumina platform (paired-end). Reads were aligned to the human reference genome/transcriptome (GRCh38/GENCODE) using STAR, and transcript-level counts were quantified using feature Counts or salmon. Differential gene expression analysis comparing dTAG-treated versus control conditions was performed using DESeq2, with genes considered differentially expressed at an adjusted p-value (Padj) < 0.05. Gene Ontology Biological Process (GO:BP) enrichment analysis of differentially expressed genes was performed using clusterProfiler (or an equivalent GO enrichment tool), with pathway enrichment expressed as the fraction of differentially expressed features relative to total pathway size and colored by Padj.

### Luciferase-based bioluminescence viability assay

Doxycycline-inducible shRNA HCT116 (p53+/+) cell lines co-expressing firefly luciferase were induced with doxycycline to knock down CIZ1 (shRNA_2) or CDC6 (shRNA_2), alongside control shRNA. Luciferase activity was measured using a luciferase substrate assay (Steady-Glo Luciferase Assay System Promega E2510) and GloMax Discover Microplate Reader system (Promega), with luminescence expressed in relative light units (RLU) as a surrogate for viable/proliferating cell number.

### Xenograft tumor studies and in vivo bioluminescence imaging

All animal procedures were performed in accordance with protocols approved by the Institutional Animal Care and Use Committee (IACUC). Doxycycline-inducible, luciferase-expressing HCT116 (p53+/+) cells stably expressing control, CIZ1 (shRNA_1), or CDC6 (shRNA_1) shRNA were harvested and resuspended in PBS/Matrigel, and injected subcutaneously into the bilateral flanks of female athymic nude mice (n = 5 mice per group, 10 tumors per group). Doxycycline was administered continuously (in drinking water and/or chow) beginning at the time of implantation to induce shRNA expression throughout the experiment. Tumor growth was monitored weekly for four weeks by intraperitoneal injection of D-luciferin (VivoGlo Luciferin, In Vivo Grade, Promega #P1042) followed by bioluminescence imaging on an IVIS imaging system (PerkinElmer). Total luminescence signal (photons/sec) was quantified for each tumor at each time point using Living Image software, with left- and right-flank tumors analyzed individually. Mice were monitored throughout for tumor burden and welfare according to approved endpoint criteria.

### Live-cell imaging and CIZ1-overexpression cytotoxicity assay

Tetracycline-inducible mGFP-CIZ1 U2OS (Flp-In T-REx) cells were induced with doxycycline and transferred to imaging dishes on the stage of a spinning-disk confocal microscope (PerkinElmer) maintained at 37°C with 5% CO2 in complete DMEM. Time-lapse images were acquired at multiple positions using a 20–40× objective at regular intervals over approximately 18–24 h to capture mGFP-CIZ1 localization dynamics and cell fate following CIZ1 induction. Images were processed and assembled into movies using ImageJ/Volocity software.

### Bioinformatic and sequence analyses

Multiple sequence alignments of ORC1 and CIZ1 across species were performed using Clustal Omega/MUSCLE and visualized with Jalview or ESPript. Intrinsic disorder prediction for human CIZ1 and mouse Ciz1 was performed using PrDOS (Prediction of Disordered regions). Domain boundaries and conserved motifs (Cy-box, CDK phosphorylation consensus S/TPxK/R, zinc-finger domains, matrin homology domain) were annotated based on Pfam/InterPro domain predictions combined with the experimentally mapped binding data presented in this study.

### Viral production and transduction

Lentiviruses were produced by co-transfecting HEK293T cells with the target plasmid and packaging plasmids psPAX2 and pVSVG (ratio 1.5:2:1) using polyethylenimine (PEI, MW 25,000; Polysciences). Medium was changed 7 h post-transfection, and viral supernatant was collected at 36 and 72 h. For pooled library screening, viral supernatant was concentrated using Lenti-X Concentrator (Takara) per the manufacturer’s protocol. Target cells were transduced by mixing with viral supernatant supplemented with 8 µg/mL polybrene and centrifuging at 1,700 rpm for 30 min at room temperature (spinfection). Fresh medium was added 24 h post-transduction, and antibiotic selection (1 µg/mL puromycin, 10 µg/mL blasticidin, 200 µg/mL hygromycin, as appropriate) was applied 72 h post-infection.

### Quantification and statistical analysis

Data are presented as mean ± SEM or mean ± SD as indicated in individual figure legends, with the number of independent biological and/or technical replicates specified in each legend. Statistical comparisons between two groups were performed using two-tailed Student’s t-test; comparisons among more than two groups were performed by one-way or two-way ANOVA with appropriate post-hoc correction, using GraphPad Prism software. A p-value < 0.05 was considered statistically significant. Immunoblot and colony-formation quantifications were performed using ImageJ software. Flow cytometry data were analyzed using FlowJo software. RNA-seq and ChIP-seq statistical analyses were performed as described in the corresponding sections above.

### Author Contributions

MH, MT and SN performed experiments, MH and BS designed the experiments and wrote the paper. BS secured funding

## REFERENCES

1. Ainscough JF-X, Rahman FA, Sercombe H, Sedo A, Gerlach B, Coverley D. 2007. C-terminal domains deliver the DNA replication factor Ciz1 to the nuclear matrix. J Cell Sci 120: 115–124.

2. Anachkova B, Djeliova V, Russev G. 2005. Nuclear matrix support of DNA replication. J Cell Biochem 96: 951–961.

3. Araki H. 2010. Cyclin-dependent kinase-dependent initiation of chromosomal DNA replication. Current opinion in cell biology 22: 766–771.

4. Attali I, Botchan MR, Berger JM. 2021. Structural Mechanisms for Replicating DNA in Eukaryotes. Annu Rev Biochem 90: 1–30.

5. Berezney R, Dubey DD, Huberman JA. 2000. Heterogeneity of eukaryotic replicons, replicon clusters, and replication foci. Chromosoma 108: 471–484.

6. Bleichert F, Botchan MR, Berger JM. 2017a. Mechanisms for initiating cellular DNA replication. Science 355: eaah6317.

7. Bleichert F, Botchan MR, Berger JM. 2017b. Mechanisms for initiating cellular DNA replication. Science 355.

8. Borlado LR, Méndez J. 2008. CDC6: from DNA replication to cell cycle checkpoints and oncogenesis. Carcinogenesis 29: 237–243.

9. Chen X, Wang P, Wang S, Li J, Ou T, Zeng X. 2019. CIZ1 knockdown suppresses the proliferation of bladder cancer cells by inducing apoptosis. Gene 719: 143946.

10. Coleman KE, Grant GD, Haggerty RA, Brantley K, Shibata E, Workman BD, Dutta A, Varma D, Purvis JE, Cook JG. 2015. Sequential replication-coupled destruction at G1/S ensures genome stability. Genes Dev 29: 1734–1746.

11. Cook PR. 1999. The Organization of Replication and Transcription. Science 284: 1790–1795.

12. Copeland NA, Sercombe HE, Ainscough JFX, Coverley D. 2010. Ciz1 cooperates with cyclin-A–CDK2 to activate mammalian DNA replication in vitro. J Cell Sci 123: 1108–1115.

13. Copeland NA, Sercombe HE, Wilson RHC, Coverley D. 2015. Cyclin-A–CDK2-mediated phosphorylation of CIZ1 blocks replisome formation and initiation of mammalian DNA replication. J Cell Sci 128: 1518–1527.

14. Costa A, Diffley JFX. 2022. The Initiation of Eukaryotic DNA Replication. Annu Rev Biochem 91: 107– 131.

15. Coverley D, Marr J, Ainscough J. 2005. Ciz1 promotes mammalian DNA replication. J Cell Sci 118: 101–112.

16. Djeliova V, Russev G, Anachkova B. 2001. Dynamics of association of origins of DNA replication with the nuclear matrix during the cell cycle. Nucleic Acids Res 29: 3181–3187.

17. Dobbs OG, Wilson RHC, Newling K, Ainscough JF-X, Coverley D. 2023. Epigenetic instability caused by absence of CIZ1 drives transformation during quiescence cycles. BMC Biol 21: 175.

18. Evrin C, Clarke P, Zech J, Lurz R, Sun J, Uhle S, Li H, Stillman B, Speck C. 2009. A double-hexameric MCM2-7 complex is loaded onto origin DNA during licensing of eukaryotic DNA replication. Proc Natl Acad Sci 106: 20240–20245.

19. Frisbie V, Bleichert F. 2026. License to Replicate: Mechanisms of Licensing Eukaryotic Origins for DNA Replication. BioEssays 48: e70095.

20. Galanti L, Pfander B. 2026. Exploring the multifaceted Dbf4-dependent kinase from temporal, spatial, and substrate repertoire perspectives. Commun Biol 9: 858.

21. Håland TW, Boye E, Stokke T, Grallert B, Syljuåsen RG. 2015. Simultaneous measurement of passage through the restriction point and MCM loading in single cells. Nucleic Acids Res 43: e150–e150.

22. Hall JR, Lee HO, Bunker BD, Dorn ES, Rogers GC, Duronio RJ, Cook JG. 2008. Cdt1 and Cdc6 Are Destabilized by Rereplication-induced DNA Damage*. J Biol Chem 283: 25356–25363.

23. Hanafiah A, Geng Z, Liu T, Tai YT, Cai W, Wang Q, Christensen N, Liu Y, Yue F, Gao Z. 2026. cPRC1.2 and CTCF-mediated transition from poised to active chromatin loops at bivalent genes. EMBO Rep 1–29.

24. He W, Zhang L, Villarreal OD, Fu R, Bedford E, Dou J, Patel AY, Bedford MT, Shi X, Chen T, et al. 2019. De novo identification of essential protein domains from CRISPR-Cas9 tiling-sgRNA knockout screens. Nat Commun 10: 4541.

25. Higgins G, Roper KM, Watson IJ, Blackhall FH, Rom WN, Pass HI, Ainscough JFX, Coverley D. 2012. Variant Ciz1 is a circulating biomarker for early-stage lung cancer. Proc Natl Acad Sci 109: E3128– E3135.

26. Hossain M, Bhalla K, Stillman B. 2021. Multiple, short protein binding motifs in ORC1 and CDC6 control the initiation of DNA replication. Mol Cell 81: 1951–1969.

27. Hossain M, Stillman B. 2012. Meier-Gorlin syndrome mutations disrupt an Orc1 CDK inhibitory domain and cause centrosome reduplication. Gene Dev 26: 1797–810.

28. Hossain M, Stillman B. 2016. Opposing roles for DNA replication initiator proteins ORC1 and CDC6 in control of Cyclin E gene transcription. eLife 5: e12785.

29. Hsu PD, Scott DA, Weinstein JA, Ran FA, Konermann S, Agarwala V, Li Y, Fine EJ, Wu X, Shalem O, et al. 2013. DNA targeting specificity of RNA-guided Cas9 nucleases. Nat Biotechnol 31: 827–832.

30. Hu Y, Stillman B. 2023. Origins of DNA replication in eukaryotes. Mol Cell 83: 352–372.

31. Igarashi T, Yano K, Endo S, Shiotani B. 2024. Tolerance of Oncogene-Induced Replication Stress: A Fuel for Genomic Instability. Cancers 16: 3507.

32. Jiang W, Wells NJ, Hunter T. 1999. Multistep regulation of DNA replication by Cdk phosphorylation of HsCdc6. Proc Natl Acad Sci 96: 6193–6198.

33. Kara N, Hossain M, Prasanth SG, Stillman B. 2015. Orc1 Binding to Mitotic Chromosomes Precedes Spatial Patterning during G1 Phase and Assembly of the Origin Recognition Complex in Human Cells*. J Biol Chem 290: 12355–12369.

34. Kotsantis P, Petermann E, Boulton SJ. 2018. Mechanisms of Oncogene-Induced Replication Stress: Jigsaw Falling into Place. Cancer Discov 8: 537–555.

35. Li N, Gao N, Zhai Y. 2023. DDK promotes DNA replication initiation: Mechanistic and structural insights. Curr Opin Struc Biol 78: 102504.

36. Li W, Xu H, Xiao T, Cong L, Love MI, Zhang F, Irizarry RA, Liu JS, Brown M, Liu XS. 2014. MAGeCK enables robust identification of essential genes from genome-scale CRISPR/Cas9 knockout screens. Genome Biol 15: 554.

37. Liu E, Lee AY-L, Chiba T, Olson E, Sun P, Wu X. 2007. The ATR-mediated S phase checkpoint prevents rereplication in mammalian cells when licensing control is disrupted. J Cell Biol 179: 643– 657.

38. Liu Q, Niu N, Wada Y, Liu J. 2016. The Role of Cdkn1A-Interacting Zinc Finger Protein 1 (CIZ1) in DNA Replication and Pathophysiology. Int J Mol Sci 17: 212.

39. Macheret M, Halazonetis TD. 2015. DNA Replication Stress as a Hallmark of Cancer. Annu Rev Pathol 10: 425–448.

40. Meers MP, Bryson TD, Henikoff JG, Henikoff S. 2019. Improved CUT&RUN chromatin profiling tools. eLife 8: e46314.

41. Mitsui K, Matsumoto A, Ohtsuka S, Ohtsubo M, Yoshimura A. 1999. Cloning and Characterization of a Novel p21Cip1/Waf1-Interacting Zinc Finger Protein, Ciz1. Biochem Bioph Res Co 264: 457–464.

42. Moreno A, Carrington JT, Albergante L, Mamun MA, Haagensen EJ, Komseli E-S, Gorgoulis VG, Newman TJ, Blow JJ. 2016. Unreplicated DNA remaining from unperturbed S phases passes through mitosis for resolution in daughter cells. Proc Natl Acad Sci 113: E5757–E5764.

43. Munkley J, Copeland NA, Moignard V, Knight JRP, Greaves E, Ramsbottom SA, Pownall ME, Southgate J, Ainscough JF-X, Coverley D. 2011. Cyclin E is recruited to the nuclear matrix during differentiation, but is not recruited in cancer cells. Nucleic Acids Res 39: 2671–2677.

44. Nasheuer HP, Meaney AM. 2024. Starting DNA Synthesis: Initiation Processes during the Replication of Chromosomal DNA in Humans. Genes 15: 360.

45. Ohta S, Tatsumi Y, Fujita M, Tsurimoto T, Obuse C. 2003. The ORC1 cycle in human cells: II. Dynamic changes in the human ORC complex during the cell cycle. The Journal of biological chemistry 278: 41535–41540.

46. Palm G, Costa A. 2025. How similar are the molecular mechanisms of yeast and metazoan genome replication initiation? Biochem Soc Trans 53.

47. Parker MW, Bell M, Mir M, Kao JA, Darzacq X, Botchan MR, Berger JM. 2019. A new class of disordered elements controls DNA replication through initiator self-assembly. eLife 8: e48562.

48. Parker MW, Botchan MR, Berger JM. 2017. Mechanisms and regulation of DNA replication initiation in eukaryotes. Crit Rev Biochem Mol 52: 107–144.

49. Pauzaite T, Thacker U, Tollitt J, Copeland NA. 2016. Emerging Roles for Ciz1 in Cell Cycle Regulation and as a Driver of Tumorigenesis. Biomol 7: 1.

50. Petersen BO, Wagener C, Marinoni F, Kramer ER, Melixetian M, Denchi EL, Gieffers C, Matteucci C, Peters J-M, Helin K. 2000. Cell cycle– and cell growth–regulated proteolysis of mammalian CDC6 is dependent on APC–CDH1. Genes Dev 14: 2330–2343.

51. Pozo PN, Cook JG. 2016. Regulation and Function of Cdt1; A Key Factor in Cell Proliferation and Genome Stability. Genes 8: 2.

52. Remus D, Beuron F, Tolun G, Griffith JD, Morris EP, Diffley JFX. 2009. Concerted Loading of Mcm2–7 Double Hexamers around DNA during DNA Replication Origin Licensing. Cell 139: 719–730.

53. Ridings-Figueroa R, Stewart ER, Nesterova TB, Coker H, Pintacuda G, Godwin J, Wilson R, Haslam A, Lilley F, Ruigrok R, et al. 2017. The nuclear matrix protein CIZ1 facilitates localization of Xist RNA to the inactive X-chromosome territory. Gene Dev 31: 876–888.

54. Rodermund L, Coker H, Oldenkamp R, Wei G, Bowness J, Rajkumar B, Nesterova T, Pinto DMS, Schermelleh L, Brockdorff N. 2021. Time-resolved structured illumination microscopy reveals key principles of Xist RNA spreading. Science 372.

55. Sofi S, Coverley D. 2023. CIZ1 in Xist seeded assemblies at the inactive X chromosome. Front Cell Dev Biol 11: 1296600.

56. Stewart ER, Turner RML, Newling K, Ridings-Figueroa R, Scott V, Ashton PD, Ainscough JFX, Coverley D. 2019. Maintenance of epigenetic landscape requires CIZ1 and is corrupted in differentiated fibroblasts in long-term culture. Nat Commun 10: 460.

57. Stillman B, Diffley JFX, Iwasa JH. 2025. Mechanisms for licensing origins of DNA replication in eukaryotic cells. Nat Struct Mol Biol 32: 1143–1153.

58. Stoynova L, Solórzano R, Collins ED. 2004. Generation of large deletion mutants from plasmid DNA. Biotechniques 36: 402–406.

59. Sugimoto N, Yoshida K, Tatsumi Y, Yugawa T, Narisawa-Saito M, Waga S, Kiyono T, Fujita M. 2009. Redundant and differential regulation of multiple licensing factors ensures prevention of re-replication in normal human cells. J Cell Sci 122: 1184–1191.

60. Swarts DRA, Stewart ER, Higgins GS, Coverley D. 2018. CIZ1-F, an alternatively spliced variant of the DNA replication protein CIZ1 with distinct expression and localisation, is overrepresented in early stage common solid tumours. Cell Cycle 60: 1–16.

61. Tarumoto Y, Lu B, Somerville TDD, Huang Y-H, Milazzo JP, Wu XS, Klingbeil O, Demerdash OE, Shi J, Vakoc CR. 2018. LKB1, Salt-Inducible Kinases, and MEF2C Are Linked Dependencies in Acute Myeloid Leukemia. Mol Cell 69: 1017–1027.e6.

62. Tatsumi Y, Ohta S, Kimura H, Tsurimoto T, Obuse C. 2003. The ORC1 Cycle in Human Cells: I. Cell Cycle-Regulated Oscillation of Human ORC1. J Biol Chem 278: 41528–41534.

63. Tennakoon C, Purbojati RW, Sung W-K. 2012. BatMis: a fast algorithm for k-mismatch mapping. Bioinformatics 28: 2122–2128.

64. Tollitt J, Preedy MK, Briggs T, Allinson SL, Staples CJ, Parsons JL, Mort RL, Copeland NA. 2026. CIZ1 regulates G1 length and the CDK threshold for initiation of DNA replication to prevent DNA replication stress. Nucleic Acids Res 54: gkag878.

65. Turvey GL, Alba EL de, Stewart E, Cook H, Alalti A, Gawne RT, Ainscough JF-X, Mason AS, Coverley D. 2025. Epigenetic deprogramming by disruption of CIZ1-RNA nuclear assemblies in early-stage breast cancers. J Cell Biol 224: e202409123.

66. Vaziri C, Saxena S, Jeon Y, Lee C, Murata K, Machida Y, Wagle N, Hwang DS, Dutta A. 2003. A p53-Dependent Checkpoint Pathway Prevents Rereplication. Mol Cell 11: 997–1008.

67. Wan H, Wang L, Huo B, Qiao Z, Zhang Y. 2023. CIZ1 aggravates gastric cancer progression via mediating FBXL19-AS1 and miR-339-3p. Heliyon 9: e21061.

68. Wang D, Wang K, Yan D, Liu J, Wang B, Li M, Wang X, Liu J, Peng Z, Li G, et al. 2013. Ciz1 is a novel predictor of survival in human colon cancer. Exp Biol Med 239: 862–870.

69. Weissmann F, Greiwe JF, Pühringer T, Eastwood EL, Couves EC, Miller TCR, Diffley JFX, Costa A. 2024. MCM double hexamer loading visualized with human proteins. Nature 636: 499–508.

70. Wells JN, Edwardes LV, Leber V, Allyjaun S, Peach M, Tomkins J, Kefala-Stavridi A, Faull SV, Aramayo R, Pestana CM, et al. 2025. Reconstitution of human DNA licensing and the structural and functional analysis of key intermediates. Nat Commun 16: 478.

71. Wilson RHC, Coverley D. 2013. Relationship between DNA replication and the nuclear matrix. Genes Cells 18: 17–31.

72. Wu J, Lei L, Gu D, Liu H, Wang S. 2016. CIZ1 is upregulated in hepatocellular carcinoma and promotes the growth and migration of the cancer cells. Tumor Biol 37: 4735–4742.

73. Yadav AK, Polasek-Sedlackova H. 2024. Quantity and quality of minichromosome maintenance protein complexes couple replication licensing to genome integrity. Commun Biol 7: 167.

74. Yang R, Hunker O, Wise M, Bleichert F. 2024. Multiple mechanisms for licensing human replication origins. Nature 636: 488–498.

75. Yin J, Wang C, Tang X, Sun H, Shao q, Yang X, Qu X. 2013. CIZ1 regulates the proliferation, cycle distribution and colony formation of RKO humQn colorectal cancer cells. Mol Med Rep 8: 1630– 1634.

76. You Z, Masai H. 2024. Assembly, Activation, and Helicase Actions of MCM2-7: Transition from Inactive MCM2-7 Double Hexamers to Active Replication Forks. Biology 13: 629.

77. Zhang D, Wang Y, Dai Y, Wang J, Suo T, Pan H, Liu H, Shen S, Liu H. 2015. CIZ1 promoted the growth and migration of gallbladder cancer cells. Tumor Biol 36: 2583–2591.

78. Zhou X, Liu Q, Wada Y, Liao L, Liu J. 2018. CDKN1A-interacting zinc finger protein 1 is a novel biomarker for lung squamous cell carcinoma. Oncol Lett 15: 183–188.

