## Supplementary material for "A role for CIZ1, a Nuclear Matrix Protein Overexpressed in Cancer Cells, in Controlling the Initiation of DNA Replication": Suplemental Figures

### Hossain-Figure S1

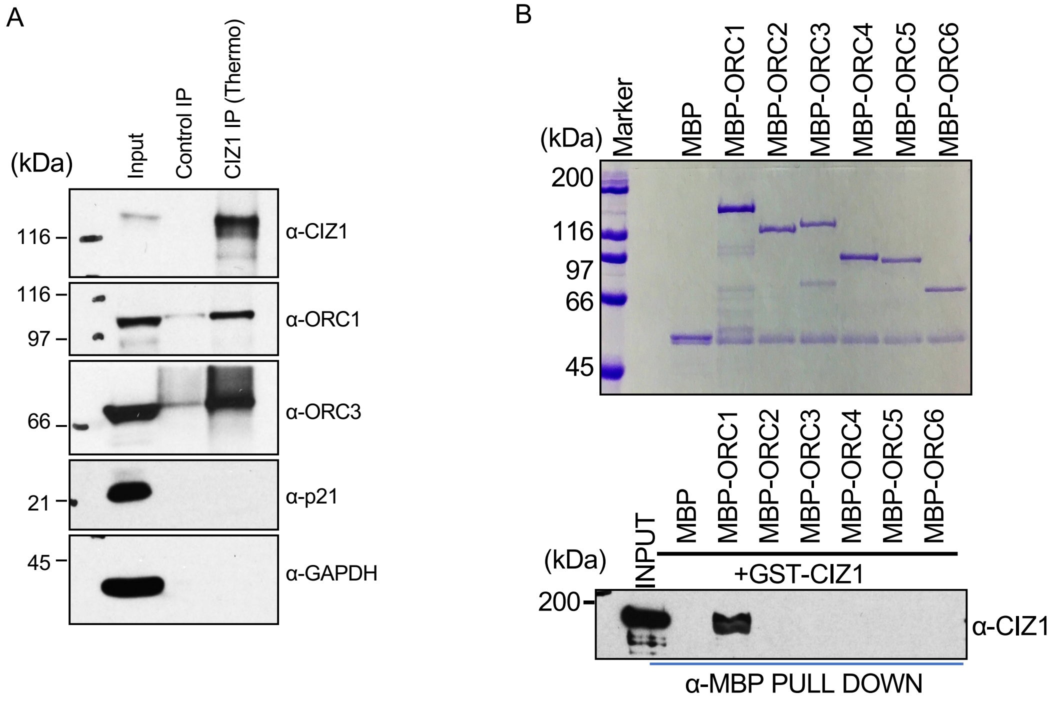


**Figure S1. CIZ1 interacts with ORC1/ORC3 and directly binds ORC subunits in vitro**

(A) CIZ1 was immunoprecipitated from asynchronously proliferating U2OS cell lysate and associated proteins ORC1 and ORC3, while no p21 interaction was observed. Input and immunoprecipitated fractions represent 1% and 10% of total lysate, respectively. Molecular weight markers are indicated in kilodaltons (kDa).

(B) Purified MBP-ORC subunit proteins bound to α-MBP antibody magnetic beads were incubated with GST-CIZ1 protein in an MBP pull-down assay and blotted with anti-CIZ1 antibody (bottom panel). The loading of MBP-ORC proteins used in pull-down assays is shown in the top panel. MBP protein served as negative control.

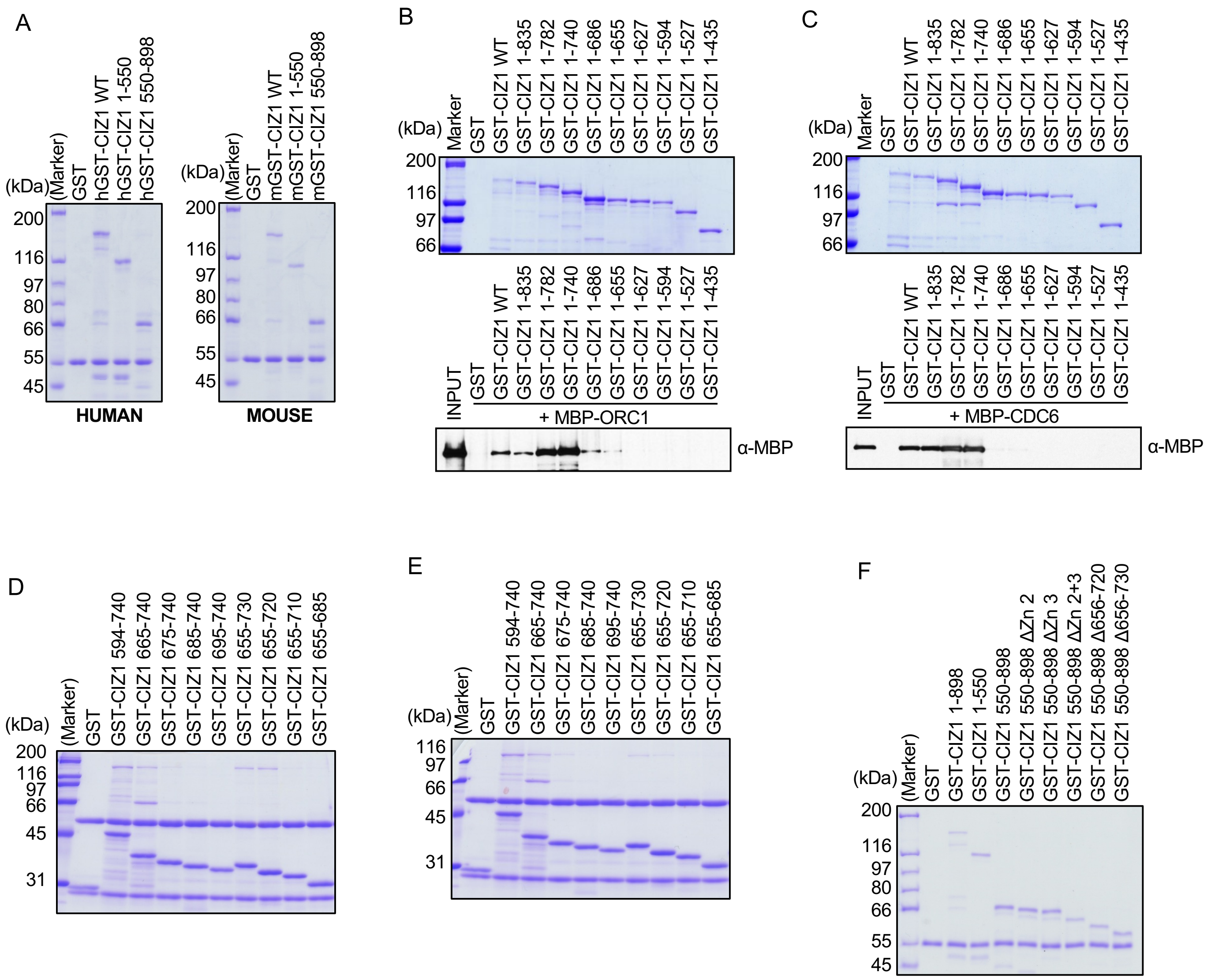


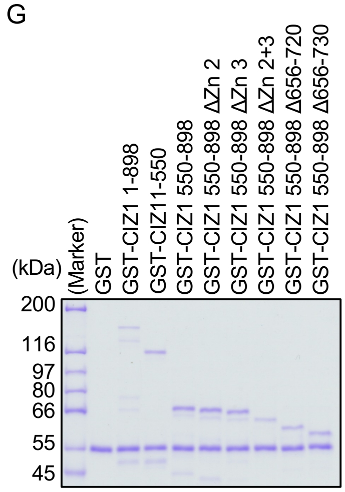


**Figure S2. Recombinant purification of GST-CIZ1 truncations and mapping of the minimal ORC1-interacting domain by in vitro GST pull-down.** (A) Coomassie-stained gels of purified GST, full-length human GST-CIZ1 WT, and human GST-CIZ1 fragments (1–550, 550–898) (left), alongside the corresponding mouse GST-CIZ1 WT and fragments (right), confirming recombinant protein purity and expected molecular weights for both species. (B & C) Coomassie-stained gel (top) of purified GST and a nested series of GST-CIZ1 C-terminal truncations (1–898/WT through 1–435), and corresponding GST pull-down immunoblot (bottom) probed with anti-MBP following incubation with MBP-ORC1 and MBP-CDC6. Robust MBP-ORC1 and MBP-CDC6 binding was retained by GST-CIZ1 1–835, 1–782, and 1–740, progressively diminished for 1–686 and 1–655, and lost for constructs truncated at or below 1–627, localizing the minimal ORC1-interacting region to approximately residues 627–740. (D & E) Coomassie-stained gel showing equitable loading of purified GST and GST-CIZ1 fragments spanning residues 594–740 with sequential C-terminal truncations (665–740 through 655–685), used to further refine the ORC1 and CDC6 binding boundary, related to Fig. 2E. (F & G) Coomassie-stained gel showing equitable loading of GST and GST-CIZ1 constructs spanning residues 550–898, including zinc-finger deletion mutants (ΔZn2, ΔZn3, ΔZn2+3) and internal deletions (Δ656–720, Δ656–730), alongside GST-CIZ1 1–898 and 1–550 used to further refine the ORC1 and CDC6 binding boundary, related to Fig. 2F.

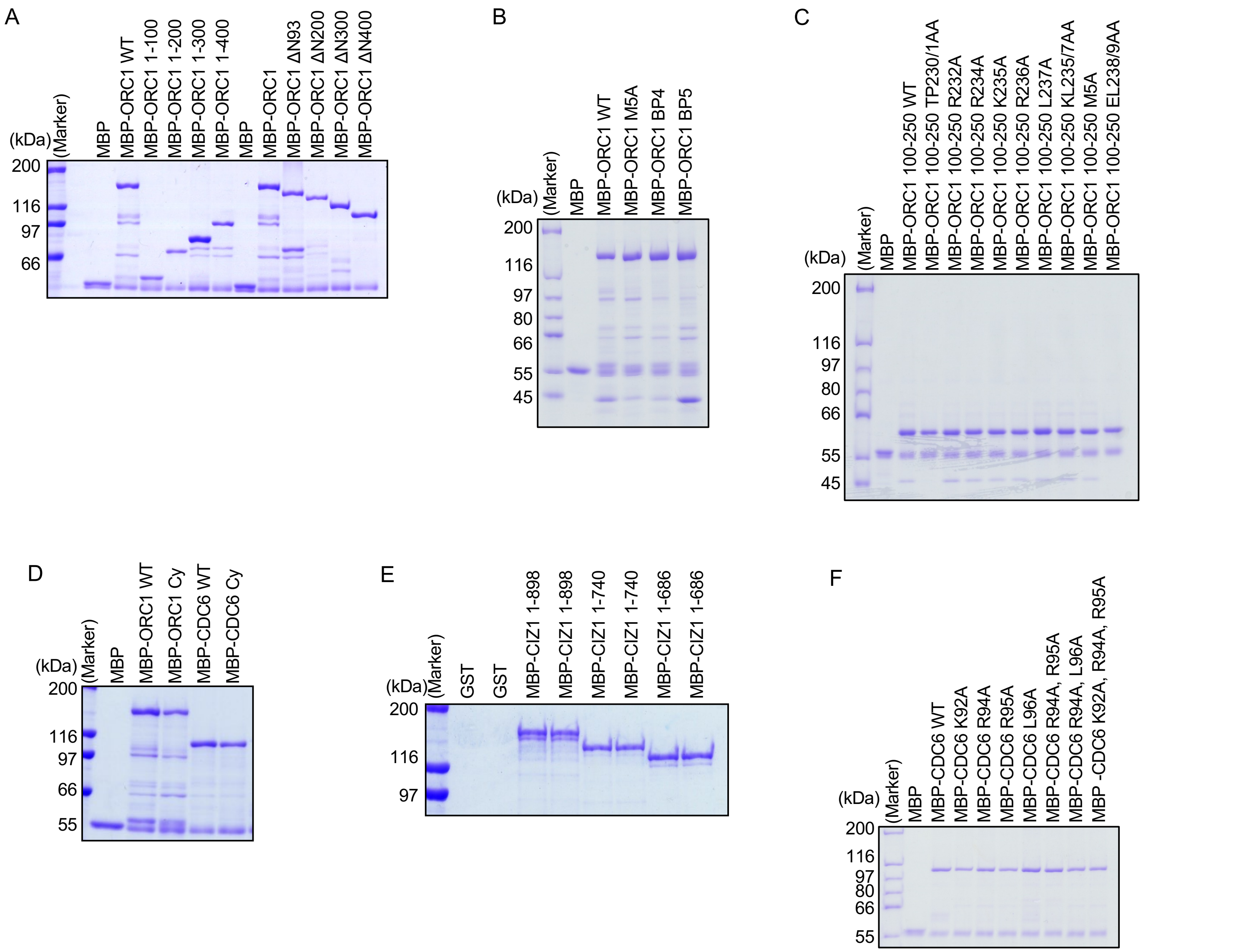


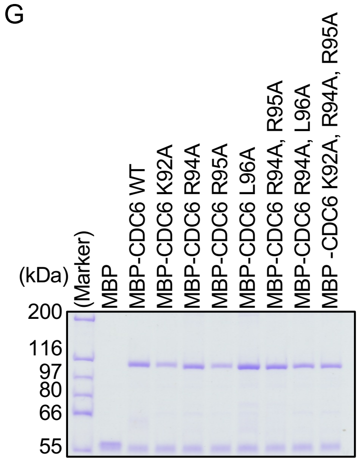


**Figure S3. CIZ1 binds ORC1 and CDC6 through overlapping N-terminal motifs that are shared with cyclin A/CDK2 binding.** (A) Coomassie-stained gel of purified MBP and MBP-ORC1 WT, N-terminal truncations (1–100 through 1–400), and N-terminal deletions (ΔN93 through ΔN400), corresponding to the pull-down inputs shown in Fig. 3A. (B) Coomassie-stained gel showing equitable loading of MBP-ORC1 WT, M5A, BP4, and BP5 combination mutants, corresponding to Fig. 3B. (C) Coomassie-stained gel showing equitable loading of MBP or MBP-ORC1 100–250 WT and point/cluster mutants used in Fig. 3C. (D) Coomassie-stained gel showing equitable loading of MBP, MBP-ORC1 WT, MBP-ORC1 Cy mutant, MBP-CDC6 WT or MBP-CDC6 Cy mutant used in Fig. 3E. (E) Coomassie-stained gel showing equitable loading of MBP, MBP-CIZ1 1–898 and 1–740 and 1–686 constructs used in Fig. 3F. (F & G) Coomassie-stained gel showing equitable loading of MBP or MBP-CDC6 WT and point mutants (K92A, R94A, R95A, L96A, R94A/R95A, R94A/L96A, K92A/R94A/R95A) used in Fig. 3G (F, incubated with GST-CIZ1 protein; G, incubated with Cyclin A-CDK2 protein).

### Hossain-Figure S4

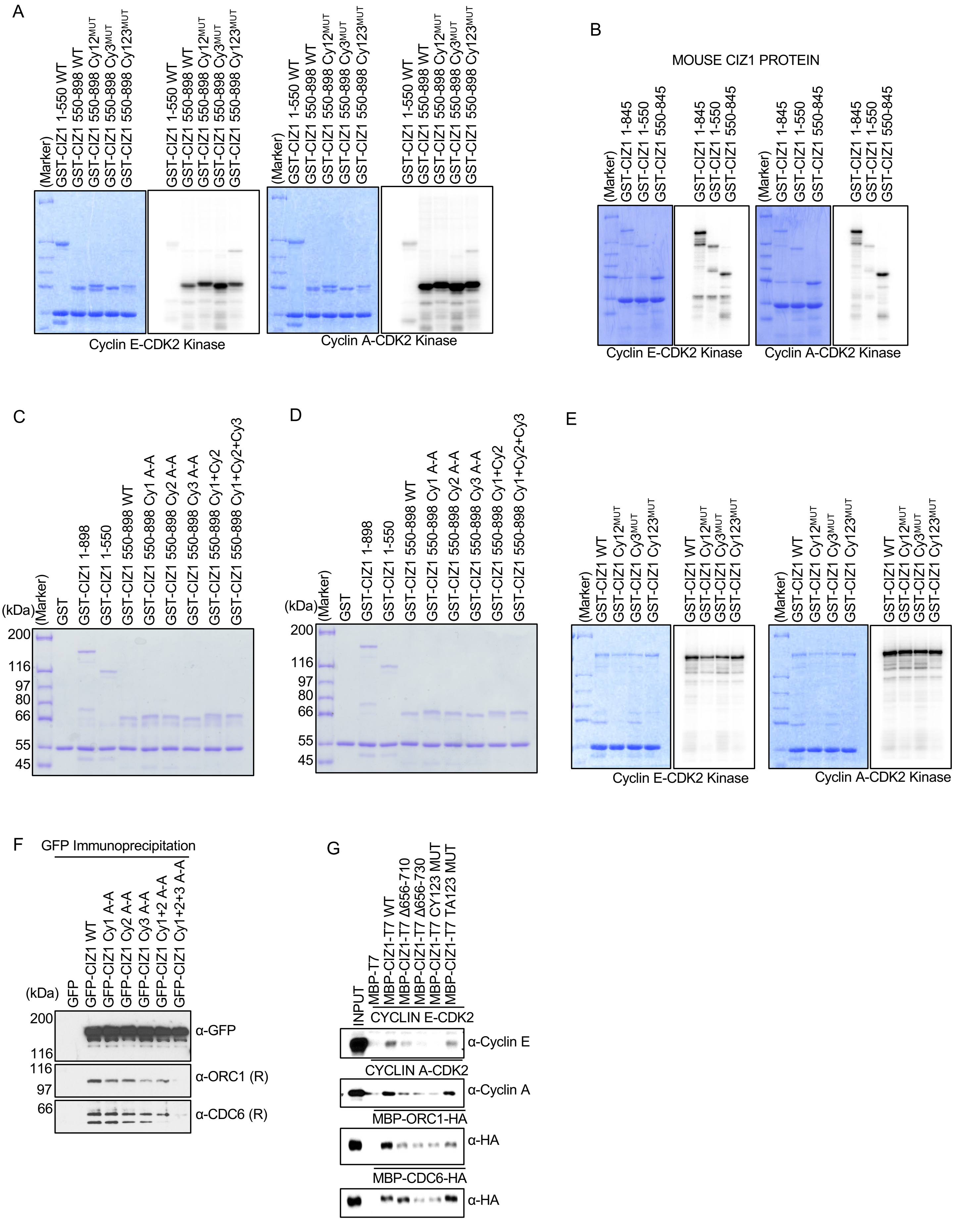

**Figure S4. Validation of CIZ1 Cy-motif and phosphorylation-site mutants, and cross-species conservation of cyclin-CDK2 binding and phosphorylation.** (A) In vitro kinase assays with Cyclin E-CDK2 (left) or Cyclin A-CDK2 (right) on GST-CIZ1 1–550 WT and 550–898 WT/Cy12$^{MUT}$/Cy3$^{MUT}$/Cy123$^{MUT}$ proteins, with Coomassie-stained input gels and corresponding phosphor-images, showing that Cy-motif mutation does not markedly impair in vitro phosphorylation by either cyclin-CDK2 complex. (B) In vitro kinase assays using mouse GST-CIZ1 1–845, 1–550, and 550–845 proteins with Cyclin E-CDK2 (left) and Cyclin A-CDK2 (right), with Coomassie-stained inputs and phosphor-images, showing conservation of cyclin-CDK2-mediated phosphorylation of the C-terminal CIZ1 region between human and mouse, but a difference in the N-terminal domain. (C) Coomassie-stained gel showing equitable loading of GST and GST-CIZ1 1–898, 1–550, and 550–898 WT/Cy-mutant constructs used in the interaction with Cyclin E-CDK2 (Fig. 4B). (D) Coomassie-stained gel of the same GST-CIZ1 construct panel used for the interaction with Cyclin A-CDK2 (Fig. 4B). (E) In vitro kinase assays with Cyclin E-CDK2 (left) and Cyclin A-CDK2 (right) comparing GST-CIZ1 WT to Cy12$^{MUT}$, Cy3$^{MUT}$, and Cy123$^{MUT}$, confirming that combined Cy-motif mutations do not substantially reduce CIZ1 phosphorylation. (F) The α-GFP immunoprecipitation from cells expressing GFP alone, GFP-CIZ1 WT, or individual/combined Cy-motif mutants (Cy1 A-A, Cy2 A-A, Cy3 A-A, Cy1+2 A-A, Cy1+2+3 A-A), immunoblotted for GFP, ORC1, and CDC6, showing that combined mutation of all three Cy motifs modestly reduces co-precipitation of endogenous ORC1 and CDC6. (G) In vitro reconstitution assay with MBP-T7 or MBP-CIZ1-T7 (WT, Δ656–710, Δ656–730, CY123 MUT, or TA123 MUT) incubated with either Cyclin E-CDK2, Cyclin A-CDK2, MBP-ORC1-HA, and MBP-CDC6-HA proteins, then immunoprecipitated with anti-T7 antibody and respectively immunoblotted for Cyclin E, Cyclin A, and HA. Deletion of CIZ1 residues 656–730 reduced association with all four binding partners, while phospho-site mutations had comparatively modest effects.

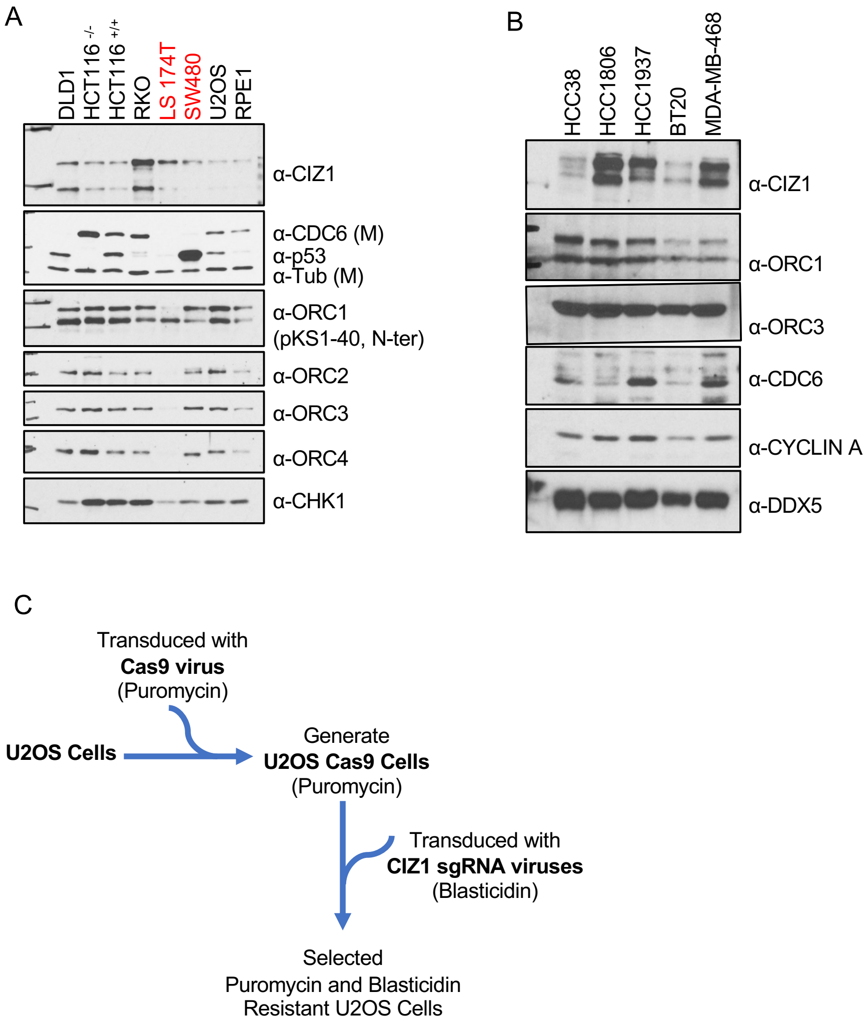


**Figure S6. CIZ1 and pre-RC component expression across additional cancer cell lines, and generation of CIZ1 CRISPR knockout cells.** (A) Immunoblot analysis of CIZ1, CDC6 (monoclonal), p53, tubulin, ORC1 (N-terminal antibody), ORC2, ORC3, ORC4, and CHK1 in DLD1, HCT116 p53-/-, HCT116 p53+/+, RKO, LS174T, SW480, U2OS, and RPE1 cell lysates, extending the panel in Figure 6A to additional colorectal lines (LS174T, SW480, highlighted in red). (B) Immunoblot analysis of CIZ1, ORC1, ORC3, CDC6, Cyclin A, and DDX5 (loading control) in a panel of triple-negative breast cancer cell lines (HCC38, HCC1806, HCC1937, BT20, MDA-MB-468), showing variable CIZ1 and pre-RC protein expression across breast cancer lines. (C) Schematic of the workflow used to generate CIZ1 CRISPR knockout U2OS cells: parental U2OS cells were transduced with Cas9 virus and selected with puromycin to generate U2OS-Cas9 cells, which were subsequently transduced with CIZ1-targeting sgRNA viruses and selected with blasticidin to yield puromycin- and blasticidin-resistant CIZ1 knockout cell populations.

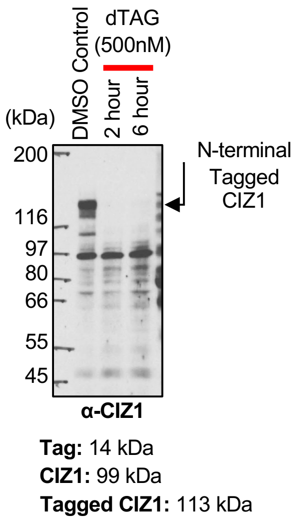


**Figure S7. Validation of dTAG-mediated degradation of FKBP12$^{F36V}$-tagged CIZ1.** Immunoblotting with anti-HA and anti-CIZ1 antibodies confirms efficient, time-dependent loss of N-terminally tagged CIZ1 (113 kDa; endogenous CIZ1, 99 kDa; tag, 14 kDa) within 2–6 h of dTAG (500 nM) treatment relative to DMSO control, validating the acute degron system used in Figure 7A.
